# Converging intracortical signatures of two separated processing timescales in human early auditory cortex

**DOI:** 10.1101/730002

**Authors:** Fabiano Baroni, Benjamin Morillon, Agnès Trébuchon, Catherine Liégeois-Chauvel, Itsaso Olasagasti, Anne-Lise Giraud

## Abstract

Neural oscillations in auditory cortex are argued to support parsing and representing speech constituents at their corresponding temporal scales. Yet, how incoming sensory information interacts with ongoing spontaneous brain activity, what features of the neuronal microcircuitry underlie spontaneous and stimulus-evoked spectral fingerprints, and what these fingerprints entail for stimulus encoding, remain largely open questions. We used a combination of human invasive electrophysiology, computational modeling and decoding techniques to assess the information encoding properties of brain activity and to relate them to a plausible underlying neuronal microarchitecture. We analyzed intracortical auditory EEG activity from 10 patients while they were listening to short sentences. Pre-stimulus neural activity in early auditory cortical regions often exhibited power spectra with a shoulder in the delta range and a small bump in the beta range. Speech decreased power in the beta range, and increased power in the delta-theta and gamma ranges. Using multivariate machine learning techniques, we assessed the spectral profile of information content for two aspects of speech processing: detection and discrimination. We obtained better phase than power information decoding, and a bimodal spectral profile of information content with better decoding at low (delta-theta) and high (gamma) frequencies than at intermediate (beta) frequencies. These experimental data were reproduced by a simple rate model made of two subnetworks with different timescales, each composed of coupled excitatory and inhibitory units, and connected via a negative feedback loop. Modeling and experimental results were similar in terms of pre-stimulus spectral profile (except for the iEEG beta bump), spectral modulations with speech, and spectral profile of information content. Altogether, we provide converging evidence from both univariate spectral analysis and decoding approaches for a dual timescale processing infrastructure in human auditory cortex, and show that it is consistent with the dynamics of a simple rate model.

**Author summary:** Like most animal vocalizations, speech results from a pseudo-rhythmic process that reflects the convergence of motor and auditory neural substrates and the natural resonance properties of the vocal apparatus towards efficient communication. Here, we leverage the excellent temporal and spatial resolution of intracranial EEG to demonstrate that neural activity in human early auditory cortical areas during speech perception exhibits a dual-scale spectral profile of power changes, with speech increasing power in low (delta-theta) and high (gamma - high-gamma) frequency ranges, while decreasing power in intermediate (alpha-beta) frequencies. Single-trial multivariate decoding also resulted in a bimodal spectral profile of information content, with better decoding at low and high frequencies than at intermediate ones. From both spectral and informational perspectives, these patterns are consistent with the activity of a relatively simple computational model comprising two reciprocally connected excitatory/inhibitory sub-networks operating at different (low and high) timescales. By combining experimental, decoding and modeling approaches, we provide consistent evidence for the existence, information coding value and underlying neuronal architecture of dual timescale processing in human auditory cortex.

## Introduction

The brains of humans and other animals generate electrical activity that often exhibits rhythmic patterns, which are apparent as shoulders or small bumps in the power spectrum on top of the 1/f^*α*^ profile (Buzsaki, 2006; Buzsáki and Draguhn, 2004). It has been suggested that rhythmic activity constitutes the neural basis of rhythmic and pseudo-rhythmic motor actions such as breathing, locomotion, chewing, peristalsis and the generation of vocalizations and other communication signals, the latter being more prominently developed in primates and birds.

Neural oscillations correspond to rhythmic activity contributed by several thousand neurons, astrocytes and possibly other cell types; they provide a compact, low-dimensional signature of the local network state that can be informative about the contextual cognitive state of the animal (e.g. its arousal (Steriade et al., 1993; McGinley et al., 2015), attentional (Besle et al., 2011; Ding and Simon, 2012; Zion Golumbic et al., 2013; Klimesch, 2012; Clayton et al., 2015; Calderone et al., 2014) and expectational (Saleh et al., 2010; Stefanics et al., 2010; Cravo et al., 2011; Rohenkohl and Nobre, 2011; Arnal and Giraud, 2012; Arnal et al., 2015; Morillon et al., 2015; Morillon and Baillet, 2017; Breska and Deouell, 2017) state) and even provide diagnostic information about pathological conditions such as schizophrenia, autism and dyslexia (Roopun et al., 2008; Lehongre et al., 2011; Lehongre et al., 2013; Calderone et al., 2014; Uhlhaas et al., 2010; Uhlhaas and Singer, 2012; Sun et al., 2013; Voytek and Knight, 2015; Soltész et al., 2013; Jochaut et al., 2015; Simon and Wallace, 2016).

Accordingly, alterations of spectral features observed in clinical populations have been related to microscopic anomalies in interneuronal function (Gonzalez-Burgos and Lewis, 2008; Pizzarelli and Cherubini, 2011) and/or in the local balance and coordination between synaptic excitation and inhibition (Fenton, 2015; Gao and Penzes, 2015). Macroscopic features related to rhythmic brain activity could hence reflect microscopic anomalies at the neuronal level and, at least in some cases, be related to specific sets of susceptibility genes (Ramamoorthi and Lin, 2011; Gao and Penzes, 2015; Benítez-Burraco and Murphy, 2016), further enhancing their interest for both basic and clinical research.

In humans, multiple pieces of evidence suggest that auditory perception, and its associated brain activity, is not a scale-free process, but presents at least two separated frequency bands, approximately located near the classically defined delta-theta (1-8 Hz) and gamma (30-60 Hz) bands, where perception and brain entrainment surpass those observed in intermediate frequencies (Poeppel, 2003; Boemio et al., 2005; Luo and Poeppel, 2012; Edwards and Chang, 2013; Ross et al., 2014; Teng et al., 2016; Teng et al., 2017). It has been suggested that the motor and the auditory system co-evolved neural mechanisms for the generation of oscillations in corresponding frequency bands, matching the natural resonance properties of the jaw, the tongue and other components of the vocal apparatus that are critical for the generation of articulated vocalizations (Morillon et al., 2010). In particular, neural oscillations in early auditory cortical regions (i.e. primary and secondary auditory cortex) are thought to be involved in the parsing and representation of speech constituents at the corresponding temporal scale, particularly at the syllabic (∼ 2-8 Hz) and phonemic (∼ 30-50 Hz) scales (Giraud and Poeppel, 2012).

In this work, we present intracranial data recorded from ten epilepsy patients while they were listening to short sentences. Spectral analysis revealed that neural activity in early auditory cortical regions in the absence of sensory stimulation does not conform to a purely scale-free process (i.e., one characterized by a 1/f^*α*^power spectrum profile): most individual subjects exhibited power spectra with a shoulder in the delta range and a small bump in the beta range. Speech stimulation decreased power in the beta range, and increased power in the delta-theta and gamma ranges.

By performing a series of spectrally-resolved decoding analyses of brain signals for speech detection and discrimination, we demonstrated that not only the information required for speech detection (i.e., distinguishing speech from pre-stimulus activity), but also information enabling speech discrimination (i.e., distinguishing between different speech segments) is preferentially conveyed in single-trial patterns of power values in low (delta - theta) and high (gamma - high-gamma) frequency bands. In some subjects, the information conveyed in single-trial patterns of phase values also exhibited such a bimodal profile whereas, at the group level, phase information was best described by a low-pass profile, with decoding accuracy decreasing with increasing frequency.

Based on our experimental observations and consistently with previous electrophysiological studies in non-human animals (Lakatos et al., 2005), we implemented a simple rate model comprised of two distinct subnetworks with different timescales, each composed of coupled excitatory and inhibitory units. The slower subnetwork exhibits a resonance in the delta-theta range (3.8 Hz), while the faster subnetwork exhibits a resonance in the gamma - high-gamma range (85.9 Hz). These frequency values roughly correspond to the delta-theta shoulder in the iEEG power spectrum in the absence of sensory stimulation, and the gamma - high-gamma bump in the iEEG power spectrum difference between speech and pre-stimulus activity, respectively; their precise values are not important since similar results were observed for a range of parameter values. They are connected via an inter-subnetwork connectivity pattern implementing a negative feedback loop between the fast and the slow subnetwork. We show that this simple model bears a remarkable resemblance to intracranial electroencephalography (iEEG) activity, both in terms of pre-stimulus spectral profile, of spectral modulations with speech, and of spectral profile of information content for speech detection and discrimination as assessed by multivariate machine learning techniques. These features do not require fine-tuning of the model parameters and are observed in a broad region of parameter space.

The current experimental findings challenge the view that extracellular field potentials in general, and electrocorticography (ECoG) and iEEG signals in particular, could be scale-free processes (Miller et al., 2009; He et al., 2010; He, 2014; Podvalny et al., 2015), and rather reveal preferred timescales in which neural responses to sensory stimuli are predominantly expressed and information about sensory inputs more effectively encoded. Our modeling investigations further show that macroscopic features of neural activity and information encoding as recorded by intracranial EEG from human early auditory cortex during speech perception can be faithfully approximated by a simple population rate model incorporating specific synaptic interactions between a slower (delta - theta) and a faster (gamma - high-gamma) neural subnetwork.

## Materials and methods

### Data Acquisition

We recorded intracranially with iEEG electrodes from 10 epilepsy patients undergoing pre-surgical monitoring. Two (six) patients were implanted with EEG electrodes on the left (right) hemisphere only, and 2 patients were implanted bilaterally. Multi-contact electrodes (0.8 mm diameter, 10 or 15 contacts of 2 mm length each with 1.5 mm spacing between contacts) were orthogonally introduced in the stereotactic space (Szikla et al., 1977; Talairach and Tournoux, 1988). Electrode location was based solely on clinical criteria. Patient age, sex, educational level, language dominance, presence of brain MRI abnormalities and locations of seizure foci are reported in Table S1. The anatomical position of each contact was then identified on the basis of (i) an axial scanner image acquired before the removal of electrodes, and (ii) an MRI scan performed after the removal of electrodes (Liegeois-Chauvel et al., 1991). Only electrodes passing through the Heschls gyrus were considered.

Intracranial EEG recordings were monopolar, with each contact of a given depth electrode referenced to an extra-dural lead using acquisition software and a 128-channel SynAmps EEG amplification system from NeuroScan Labs (Neurosoft Inc.). During the acquisition, the iEEG signal was high-pass filtered at 0.5 Hz and amplified with an anti-aliasing filter at 200 Hz (temporal resolution of 1 ms and amplitude resolution of 1 *µ*V). Sampling rate for the iEEG signal was 1000 Hz.

We did not record for 12 hours after any generalized seizure event. Voltage traces from each electrode and each trial were visually inspected; one trial from subject S8 included large amplitude high-frequency deflections of clearly non-physiological origin, and was thus excluded from the analyses. Intracranial recordings exhibit very little contamination from non-neural (e.g., muscular) sources (with the exception of specific regions of the temporal pole that can exhibit activity related to eye movements (Jerbi et al., 2009; Kovach et al., 2011)). Isolated interictal epileptiform discharges could have occurred; however, their timing, frequency and amplitude are expected to be unrelated to our variables of interest.

### Experimental protocols

In this study we considered data from 2 different auditory protocols: a functional localizer and a speech (phrase) protocol.

In the functional localizer, 30 ms pure tone sounds with 0.3 ms on and off ramps were presented binaurally (except for subject S7, who received monaural stimuli). The interstimulus interval was uniformly distributed in the interval 1030 ms + [-200,200] ms for the bilaterally implanted patients, while it was constant and fixed at 1030 ms for the other patients. Pure tone frequencies varied across subjects and are reported in Table S2 together with the number of trials in each protocol.

In the phrase protocol, subjects listened to several repetitions of two 2.5 s long sentences in French, uttered by a French female whose voice had a fundamental frequency of 201 Hz. Both trial types correspond to the French sentence “le nouveau garde la porte”, pronounced with different prosody: in the first trial type (hereafter P1), a short pause occurred after “garde”, while in the second trial type (hereafter P2), a short pause occurred after “nouveau”. Stimuli were presented monaurally in a pseudo-randomized order which was the same for every subject (P1, P1, P2, P2, P2, …) with an interstimulus interval of 4135 ms, and only the (dominant) contra-lateral response was taken into account (as the auditory ascending pathway is partly decussated). In both protocols, stimuli were delivered through headphones in a pseudo-randomized order at a 22 kHz rate using E-prime 1.1 (Psychology Software Tools Inc., Pittsburgh, PA, USA). Patients were instructed to passively listen and concentrate on what they heard. A subset of this dataset has previously been reported in (Morillon et al., 2012; Fontolan et al., 2014).

### Data analysis

#### Data preprocessing

We bipolar re-referenced the original iEEG signals to remove low spatial frequency components and hence obtain a more localized signal to better exploit the fine spatial resolution of iEEG recordings.

#### ERP analysis and channel selection

We measured Event-Related Potentials (ERP) to the pure tone stimuli to functionally select the channels that recorded activity from early auditory cortical areas (i.e. primary and secondary auditory cortex, corresponding to Brodmann areas 41 and 42). The raw voltage traces were aligned with respect to tone onset and averaged across trials for each frequency value. ERP amplitude was defined as the difference between the maximum and the minimum value in the time interval [0,200] ms post-tone onset, expressed in units of baseline standard deviation (SD). Baseline SD was measured by calculating the standard deviation across trials for each time point, and then averaging across time points. For each channel, the maximum ERP amplitude across the tested frequencies was defined as the ERP amplitude for that channel. For each multi-contact electrode, only the channel with the largest ERP amplitude was retained for further analysis (one channel per patient for the unilaterally implanted patients, two channels for the bilaterally implanted patients). The data corresponding to right ear stimulus presentation for one of the bilateral patients was lost, and the selected channel corresponding to right ear stimulus presentation for the other bilateral patient was discarded because of non-significant ERP size (1.57 SD), resulting in a total number of 10 channels, one for each subject, that entered the main analysis. ERP amplitudes for these channels were in the interval [4,18] SD. Spectral tuning curves at non-selected channels were the same as those at the selected channel for each multi-contact electrode, but exhibited ERP of lesser amplitude (i.e., lower signal-to-noise ratio); hence, larger inter-trial variability. This indicates that our iEEG multi-contact electrodes were not sampling from cortical regions with different favorite frequency, at least within the frequency resolution tested.

#### Spectral analyses

We considered spectrotemporal representations of iEEG or simulated signals based on the Continuous Wavelet Transform (CWT), using a complex Morlet wavelet with bandwidth parameter equal 1.5. To improve visualization and yield a distribution that is closer to normal, we applied a non-linearity (10log_10_) to the power values. For the estimation of phase consistency across trials, we calculated the Phase Locking Value (PLV) (Tallon-Baudry et al., 1996; Lachaux et al., 1999):

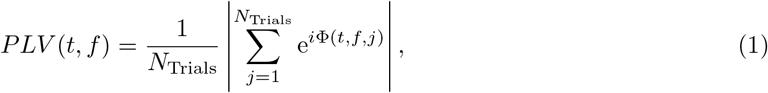

where *N*_Trials_ is the number of trials and Φ(*t, f, j*) is the phase angle in radiants for the considered time point, frequency value and trial.

To quantify the average power spectra during the pre-stimulus baseline and during speech stimulation, we estimated spectral power separately for the baseline ([-1000, 0] ms) and the speech period ([0, 1000] ms) by computing CWT-transformed signals for each time window separately, extracting 10log_10_ power, and then averaging over trials and time points within each time window. Voltage traces were normalized in each trial before CWT transformation by removing the mean value in each time window, and dividing by the standard deviation of the pre-stimulus baseline window.

For the estimation of phase consistency across trials during the pre-stimulus baseline and during speech stimulation, phase was extracted from the CWT-transformed signals calculated separately for the baseline ([-1000, 0] ms) and the speech period ([0, 1000] ms) for each trial, each frequency value and each time point. Then, the PLV value for each time window was calculated as

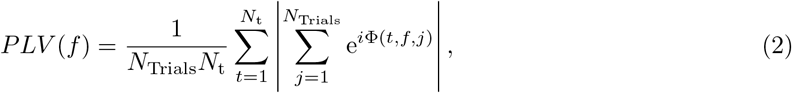

where *N*_t_ is the number of time points.

### Model

We implemented a neuronal network composed of rate units of the Wilson-Cowan type (Wilson and Cowan, 1972), adapted from (Mejias et al., 2016).

The network is composed of two subnetworks: a fast (G) subnetwork and a slow (T) subnetwork, exhibiting activity in the gamma - high-gamma and delta-theta range, respectively. Each subnetwork comprises an excitatory and an inhibitory population, mutually connected, whose average firing rates evolve according to the following equations:

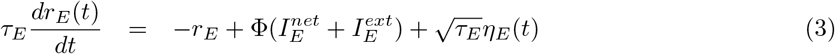

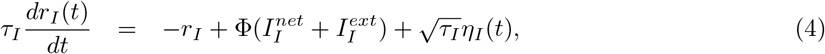

where *r*_E(I)_ is a dynamical variable describing the (dimensionless) average firing rate of the excitatory (inhibitory) population, and *τ*_E(I)_ is the corresponding time constant. The input term *I*^*net*^ represents the synaptic inputs arriving from other populations in the network, while the input term *I*^*ext*^ represents the inputs from external sources, such as sensory stimuli or other areas not explicitly included in the model. Input terms are passed through a rectifying non-linearity Φ(*x*) = *x*/(1 − *e*^−*x*^). Taking into account only local, intra-subnetwork, contributions, the network input is given by

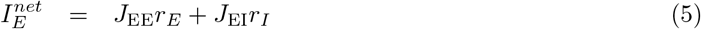

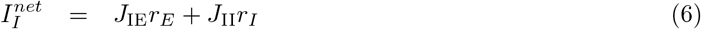

In the linear approximation (which is perfectly realized when *I*^*ext*^ ≫ *I*^*net*^) and ignoring the inter-subnetwork coupling, each subnetwork is characterized by a pair of complex conjugate eigenvalues with negative real part (indicating stable dynamics), and imaginary parts corresponding to a natural frequency of 85.9 Hz for the fast subnetwork, and of 3.8 Hz for the slow subnetwork.

Each population also receives a noise term *η*(*t*), which represents intrinsic and synaptic noise and other sources of variability. It is implemented as colored noise (i.e., a Ornstein-Uhlenbeck process), which evolves according to the equation

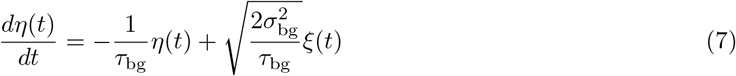

where *τ*_bg_ is the auto-correlation time constant, and *σ*_bg_ is the standard deviation. In the canonical model used throughout this study, the only inter-subnetwork connections are those from Ge to Te, and from Te to Gi. This inter-subnetwork connectivity scheme implements a negative feedback loop from the fast to the slow subnetwork (from Ge to Te), and back to the fast subnetwork (from Te to Gi). Alternative inter-subnetwork connectivity schemes have also been explored. However, they resulted in poorer fits to the iEEG dataset, hence they are not reported here.

To deliver speech input to the model, we first extracted the envelope of the speech signal. We used a time-domain method implemented in Matlab (the “peak” method for the function envelope.m), which estimates the upper and lower envelopes of a signal by using spline interpolation of the maxima and minima of the signal, respectively. After extraction of the upper and lower envelopes, we took the maximum of the absolute value of the two in each time point as an estimate of the total envelope *I*_se_(*t*). Speech input to the network is mediated by short-term synaptic depression, as often observed in thalamocortical synapses (Thomson and Deuchars, 1994; Elhilali et al., 2004). Short-term synaptic depression has been implemented according to the Tsodyks-Markram model (Tsodyks et al., 1998): we considered synaptic transmission to be mediated by a dynamical variable *x* representing the average fraction of available neurotransmitters in the bottom-up pathway conveying the speech input, which evolves as

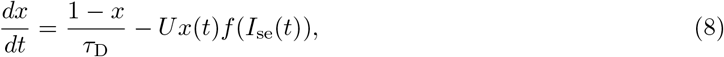

where f is the sigmoid function 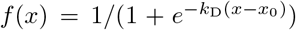. The expression above describes how the fraction of available neurotransmitters decays with incoming sensory input with rate U, and it recovers with time constant *τ*_D_. Finally, the speech input to the network, delivered to the population Ge, is calculated as follows:

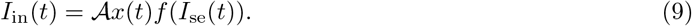

Parameter values and descriptions are provided in Table S3.

### Multivariate classification analyses

We estimated the amount of information conveyed by neural signals using binary Regularized Least-Square Classifiers (RLSC, (Rifkin et al., 2003)) with regularization parameter *λ* = 10^6^. Regularized Least-Square Classification is a machine learning technique that estimates the linear separability between patterns according to their class.

Our goal was to assess the amount of information conveyed by a spectrotemporal representation of the iEEG signal in each trial, obtained by applying a complex Morlet wavelet transformation, about the presented acoustic stimulus. In particular, we considered either 10log_10_ power, or phase angle (after taking sine and cosine), or both, at 12 × 17 (time × frequency) points for each time window in each trial, sampled from a grid with temporal duration equal 360 ms, as the input to the classifiers. Sampled points in time were evenly spaced, while points in frequency were sampled with higher resolution for lower frequencies (4 Hz intervals from 4 to 20 Hz, 10 Hz interval from 20 to 80 Hz, 20 Hz interval from 80 to 200 Hz).

This approach is similar to previous work that used multivariate patterns obtained from multi-taper spectral estimates (Tsuchiya et al., 2008; Baroni et al., 2017), but here we considered wavelet-based spectral estimates to leverage their excellent time-frequency resolution especially at intermediate and high frequencies, which could be beneficial given the highly dynamic nature of speech stimuli, and the fact that beta and gamma events tend to occur in short-lived bursts, as observed in our dataset and previously reported by others (e.g., (Sherman et al., 2016; Lundqvist et al., 2016)).

We also performed a set of spectrally-resolved decoding analyses where either power or phase patterns were sampled across time for a set of frequency values, unevenly spaced as described above, hence resulting in 12-dimensional patterns as inputs to the classifiers.

A set of weights that optimally separate trials according to their class is determined using a subset of the available trials, denoted as training set. The performance of the classifier is defined using a different set of trials, denoted as test set, as the area under the Receiver Operating Characteristic (ROC) curve, which we refer to as A’ (A prime). We report the average A’ values over *N*_iter_ cross-validations. In each cross-validation, we randomly chose a set of 0.7 × min(*N*_1_, *N*_2_) (rounded to the nearest integer) trials of each class as the training set, where *N*_1_ and *N*_2_ are the number of trials in class 1 and 2, respectively. As the test set, we chose min(*N*_1_, *N*_2_) − round(0.7 × min(*N*_1_, *N*_2_)) trials of each class among those that are not already included in the training set. Before being fed to the classifier, inputs were z-transformed: the mean and standard deviation of the relevant variable at each time-frequency point in the training set was calculated, and used to transform both training and test sets. Then, optimal RLSC weights were estimated using training trials, and their capacity to separate test trials according to their class was measured as the area under the ROC curve (A’). The number of cross-validations *N*_iter_ was set to 1000 for all the decoding analyses.

Significance of A’ values was estimated via a permutation-based statistics. For each classification considered, the class labels were randomly shuffled. Then, the average A’ value over *N*_iter_ realizations of training and test sets was calculated as described above. This procedure was repeated *N*_perm_=1000 times, yielding a probability distribution of average A’ values corresponding to the null hypothesis of lack of linear separability between the two classes. An empirical average A’ value was considered significant at level p if it exceeded the p-percentile of the corresponding null distribution (p=0.01, p=0.001). Significance thresholds at p=0.01 were estimated separately for each classification considered. In order to improve the estimation of the significance threshold at p=0.001, null A’ values were pooled across decoding types, and the corresponding significance threshold was calculated from the resulting null distribution.

We considered 3 distinct time windows for each sentence: a baseline time window ([-680 −320] ms with respect to speech onset), a speech onset time window, corresponding to the utterance of the speech token “le nou” (hereafter T1; [0 360] ms from speech onset), and a later speech time window, corresponding to the utterance of the speech token “la po” (hereafter T2; [1704 2064] ms for P1, [1589 1949] ms for P2). We considered all possible within-phrase binary decoding (T1 vs. baseline, T2 vs. baseline, and T1 vs. T2, for both P1 and P2), as well as the binary decoding analyses between corresponding speech tokens in P1 and P2.

To simplify the presentation of our results, we further grouped these decoding analyses into three categories: detection (speech token vs. baseline, comprising 4 decoding analyses: T1 vs. baseline and T2 vs. baseline for both P1 and P2), easy discrimination (between different speech tokens, or the same speech token immediately following continuous speech or occurring after a pause, comprising 3 decoding analyses: T1P1 vs. T2P1, T1P2 vs. T2P2, and T2P1 vs. T2P2), and hard discrimination (between the same speech token occurring at sentence onset in the two phrases: T1P1 vs. T1P2).

In spite of its stochastic nature, the rate model presented in the previous subsection does not fully capture the extent and complexity of the inter-trial variability observed in the iEEG dataset; in fact, the rate model yields perfect decoding accuracy in all the detection and discrimination decoding analyses just described. Hence, in order to probe the encoding properties of the model with stimuli that are similar to those employed in the iEEG experiments, we generated a set of stimuli by interpolating between P1 and P2 for each time point: 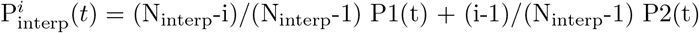, with i=1, 2, … N_interp_ and N_interp_=10.

To assess the degree of specificity vs. generalization in the read-out of the information conveyed by patterns of either power or phase values, we performed a series of decoding analyses where training and test patterns were extracted from different frequency values. This approach resembles the temporal generalization method (King and Dehaene, 2014), but generalization is assessed across the frequency instead of the time dimension.

## Results

### Spectral analysis of neural responses to speech

Most selected channels from early auditory cortex exhibited trajectories with marked and reproducible deflections in correspondence to certain acoustic landmarks, most prominently sentence onset and speech tokens that follow a pause (Fig. 1B). Another distinctive feature was the presence of a power spectral peak or a shoulder in the beta range, which decreased in amplitude with speech presentation. Speech stimulation also resulted in power increases in low (delta-theta) and high (gamma - high-gamma) frequency ranges.

**Figure 1.**
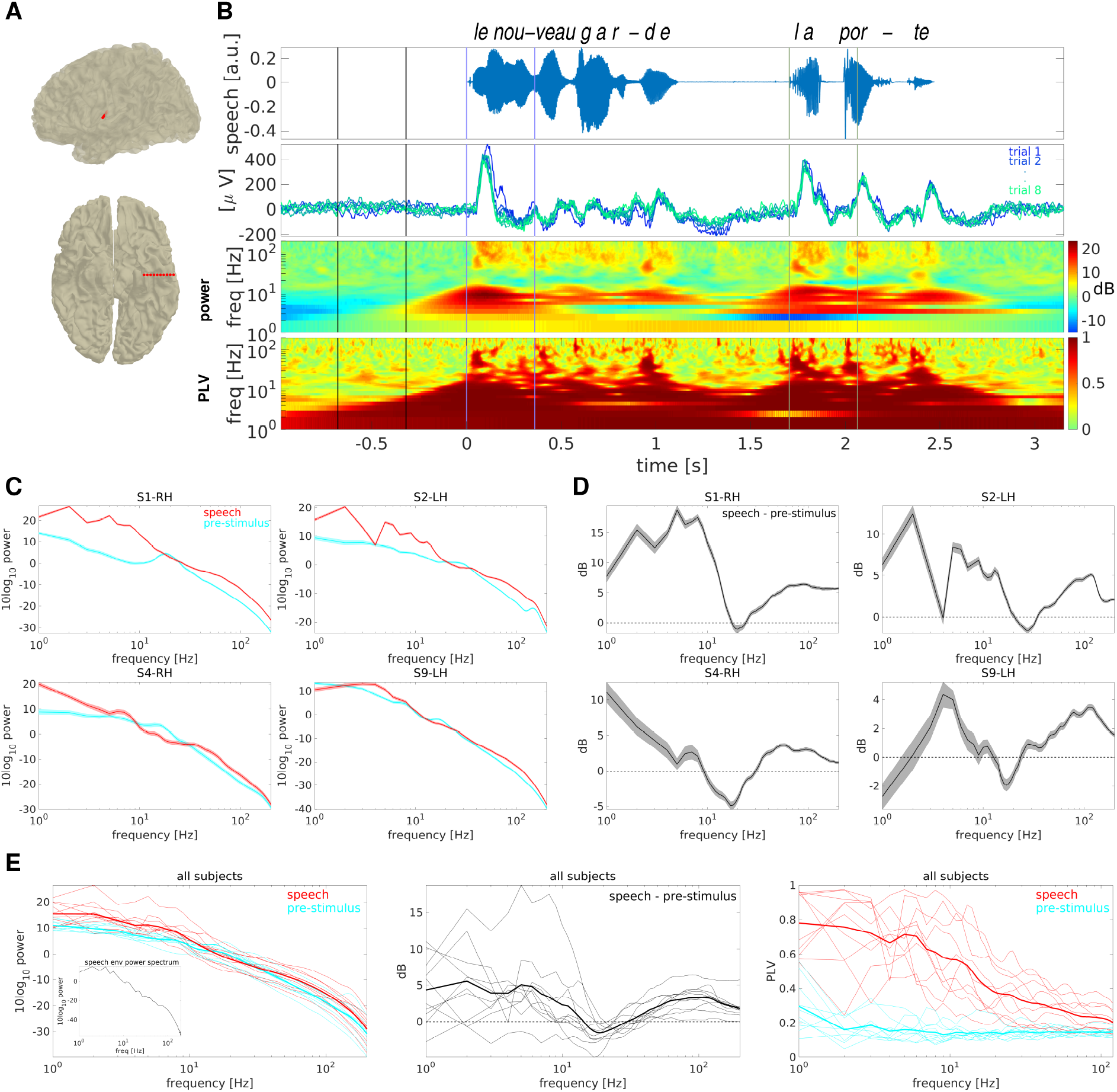
Neural responses to speech in early cortical areas. A: Anatomical image showing the multi-contact electrode implantation in early auditory cortex in an example subject on a standard brain. B: Stimulus and neural responses recorded from a strongly speech responsive channel from subject S1. Top: input stimulus corresponding to the French sentence “le nouveau garde la porte”. Middle-top: voltage trajectories for the first 8 trials. Middle-bottom: trial-averaged wavelet power spectrogram. Differences in 10log_10_ power with respect to a pre-stimulus baseline ([-1000,0] ms) are shown for the ease of visualization. Bottom: PLV spectrogram. Vertical lines indicate the three 200 ms time windows used for the speech detection and discrimination analyses shown in subsection “Stimulus encoding occurs preferentially at delta-theta and gamma frequencies”, which corresponds to pre-stimulus activity (black), “le nou” (purple) and “la po” (green). Corresponding panels for all selected channels and for both phrases are shown in Fig. S1. C: Trial-averaged power spectra calculated separately for the pre-stimulus (cyan) and the speech (red) period for four representative subjects. Shaded areas indicate s.e.m. Corresponding panels for all selected channels are shown in Fig. S2. D: As in C), but the difference in spectral power between the speech and the pre-stimulus period is shown. Corresponding panels for all selected channels are shown in Fig. S3. E: Left, middle: As in C), D), for each selected channel (thin lines). Average quantities over the set of selected channels are shown with thick lines. Inset shows the power spectrum of the speech envelope signal. Right: wavelet PLV, calculated separately for the pre-stimulus (cyan) and the speech (red) period, for each selected channel (thin lines) and averaged over the set of selected channels (thick lines). Phase-locking values for each selected channel are shown separately in Fig. S4.

Every subject exhibited this bimodal profile of spectral power changes with speech stimulation; however, the frequency ranges where power increased with stimulation and those where power decreased with stimulation exhibited some inter-individual variability (Fig. 1C-E; see also Fig. 2F). In particular, the transition between the low-frequency activation and the mid-frequency deactivation could vary between 7 and 22 Hz, and the transition between the mid-frequency deactivation and the high-frequency activation could vary between 20 and 40 Hz.

**Figure 2.**
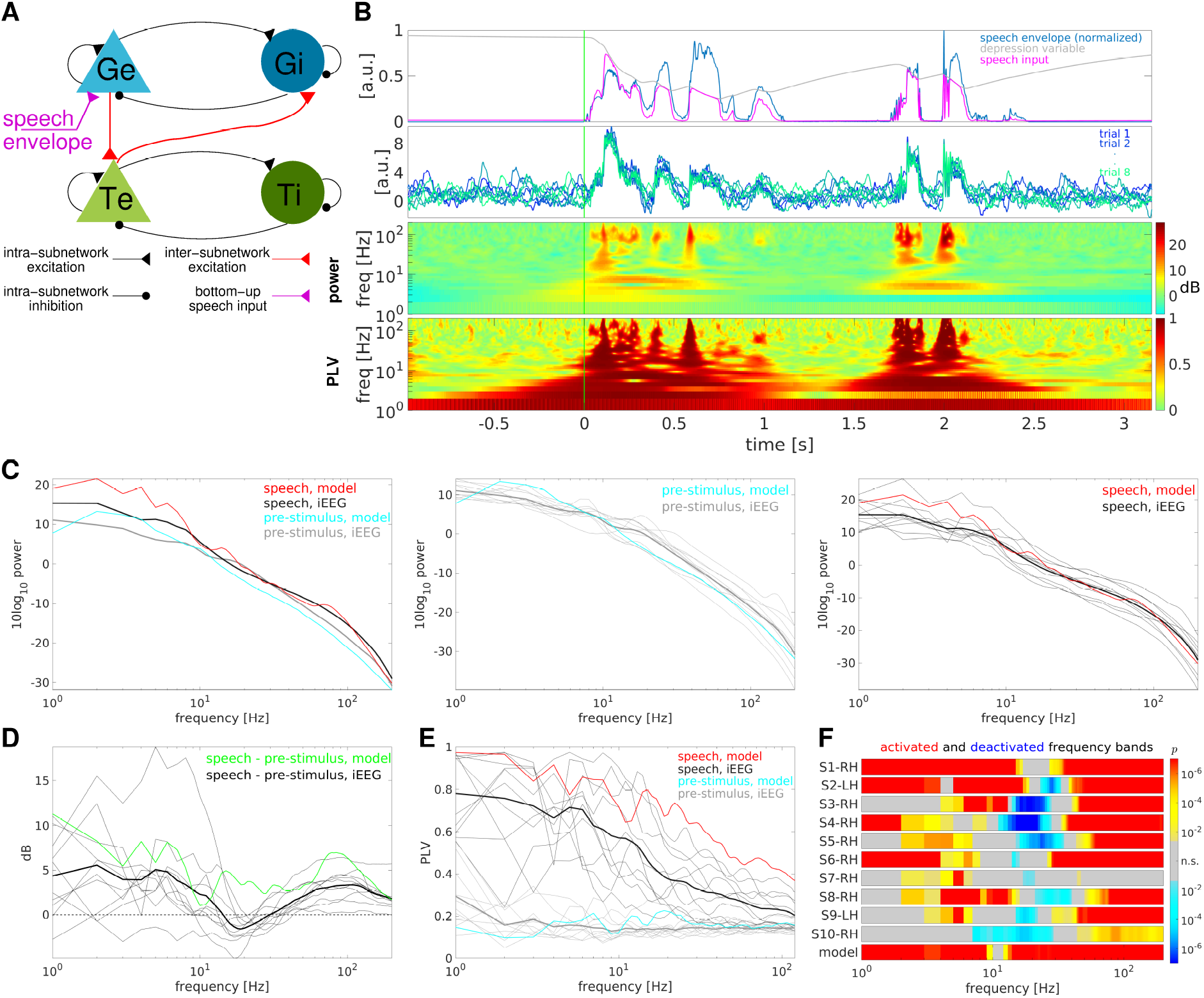
Responses to speech in a population rate model. A: Model diagram. Shades of blue [green] indicate units belonging to the fast (gamma) [slow (theta)] subnetwork. Excitatory (inhibitory) units are indicated by triangles (circles). Excitatory (inhibitory) connections are shown as lines ending in triangles (circles). Black (red) lines indicate intra- (inter-) subnetwork connections, while the magenta line indicates the bottom-up sensory input. B: The model responses to speech are shown in the same format as in Fig. 1B, with the exception of the top subpanel, which illustrates how synaptic short-term depression is implemented in the model, showing the scaled speech envelope (blue line), the dynamical variable representing the fraction of available neurotransmitters (green), and the signal injected to Ge units (magenta), resulting from the product of the first two quantities for each time point. Synaptic depression qualitatively reproduces the large responses to speech onsets and attenuated responses to subsequent syllables observed in the iEEG recordings. C: Trial-averaged power spectra calculated separately for the pre-stimulus and the speech period are shown in the left panel for the model (in cyan and red, respectively) and for the iEEG sample (average across selected channels; shown in gray and black, respectively). Power spectra corresponding to the pre-stimulus (speech) period only are shown for the model and each selected iEEG channel in the middle (right) panel. D: As in Fig. 1E, middle panel, with the addition of the corresponding model result (green line). E: As in Fig. 1E, right panel, with the addition of the corresponding model results (cyan for the pre-stimulus period, red for the speech period). F: Activated and de-activated frequency bands. For each subject and for each frequency value, the p-value from a one-tailed t-test between the speech and the baseline power spectra is shown (results from right-tail [left-tail] tests, which indicate higher power during speech [baseline], are shown in red [blue]). Only p-values smaller than p_crit_ (corresponding to a FDR q value of 0.05) are shown, and only if at least two adjacent frequency values show concordant effects.

In spite of inter-individual variability in these frequency bands, the average power spectra across the population of selected channels exhibited features that are consistent with the single-channel analyses: the presence of a beta peak during baseline, which decreases with stimulation, and a bimodal profile of activation in low (delta-theta) and high (gamma - high-gamma) frequencies, separated by an intermediate deactivated frequency band.

The pattern of phase-locking was also variable across subjects; however, most subjects as well as the mean profile across subjects exhibited a low-pass profile with strong phase-locking in the delta-theta range, and progressively lower phase-locking at higher frequencies (Fig. 1E, right panel). It is worth noting, however, that some individual subjects exhibited a band-pass PLV profile, with a peak ~ 2 Hz. Given previous work it is likely that a band-pass PLV profile with a peak ~ 1-2 Hz could have been identified in all subjects if longer continuous speech segments had been used, allowing for the assessment of phase-locking at lower frequencies (Edwards and Chang, 2013). As our main interest lies in the characterization of neural processes that could potentially underlie speech perception at the (sub-)phonemic, syllabic and word-level scales, frequencies below 1 Hz were not investigated.

### A population rate model reproduces the spectral features observed in the iEEG data

We assessed the extent to which a simple population rate model is capable of reproducing the spectral features of neural responses presented in the previous section. To this end, we implemented a population rate model following the formalism initially proposed by Wilson and Cowan (Wilson and Cowan, 1972). The model comprises two subnetworks, each composed by an excitatory and inhibitory unit with self and reciprocal connections. Further details, including the model equations, are presented in subsection “Model”.

In spite of its simplicity, the population rate model can qualitatively reproduce several features of iEEG recordings. In particular, the power spectrum during baseline exhibits a knee in the delta-theta range, and an approximately 1/f^*α*^ profile at higher frequencies. However, it lacks a beta peak, which was observed in all subjects. In response to speech, spectral power increases in low (delta-theta) and high (gamma - high-gamma) frequencies, with no significant change in the intermediate (alpha-beta) range. This profile is similar to what observed in the iEEG data. However, while all iEEG subjects presented a significant beta power reduction with speech, the model does not exhibit significant power changes in this frequency range (Fig. 2F).

### The rate model fits the iEEG data to a degree that is comparable to intersubject variability

To quantify the similarity between model and iEEG trajectories, we conducted a spectrally-resolved comparison between the model and each iEEG subject, and an analogous spectrally-resolved comparison between each pair of iEEG subjects. The similarity between the model and each iEEG subject was assessed by computing, for each subject and for each frequency value between 1 and 200 Hz, the cross-correlation between the 10log_10_ power trajectories for each pair of trials 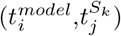, where 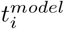 is a trial from the model simulation and 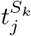 is a trial from the iEEG subject S_*k*_. Then, cross-correlations were averaged across trail pairs.

Our goal here was not to assess whether the model tends to lead or lag the iEEG data, but rather to assess the similarity between model and iEEG activity at the single-trial level, and to relate it to the degree of similarity between iEEG subjects, in a spectrally-resolved fashion. Hence, we also considered the opposite ordering, that is, the trial-averaged cross-correlation between each iEEG subject and the model (obtained from the average cross-correlation between the model and each iEEG subject by reversing the time delay axis), and averaged across orderings. Then, the maximum cross-correlation was retained for each subject and frequency value, and was then averaged across subjects to obtain a spectrally-resolved profile of similarity between model and iEEG (Fig. 3A).

**Figure 3.**
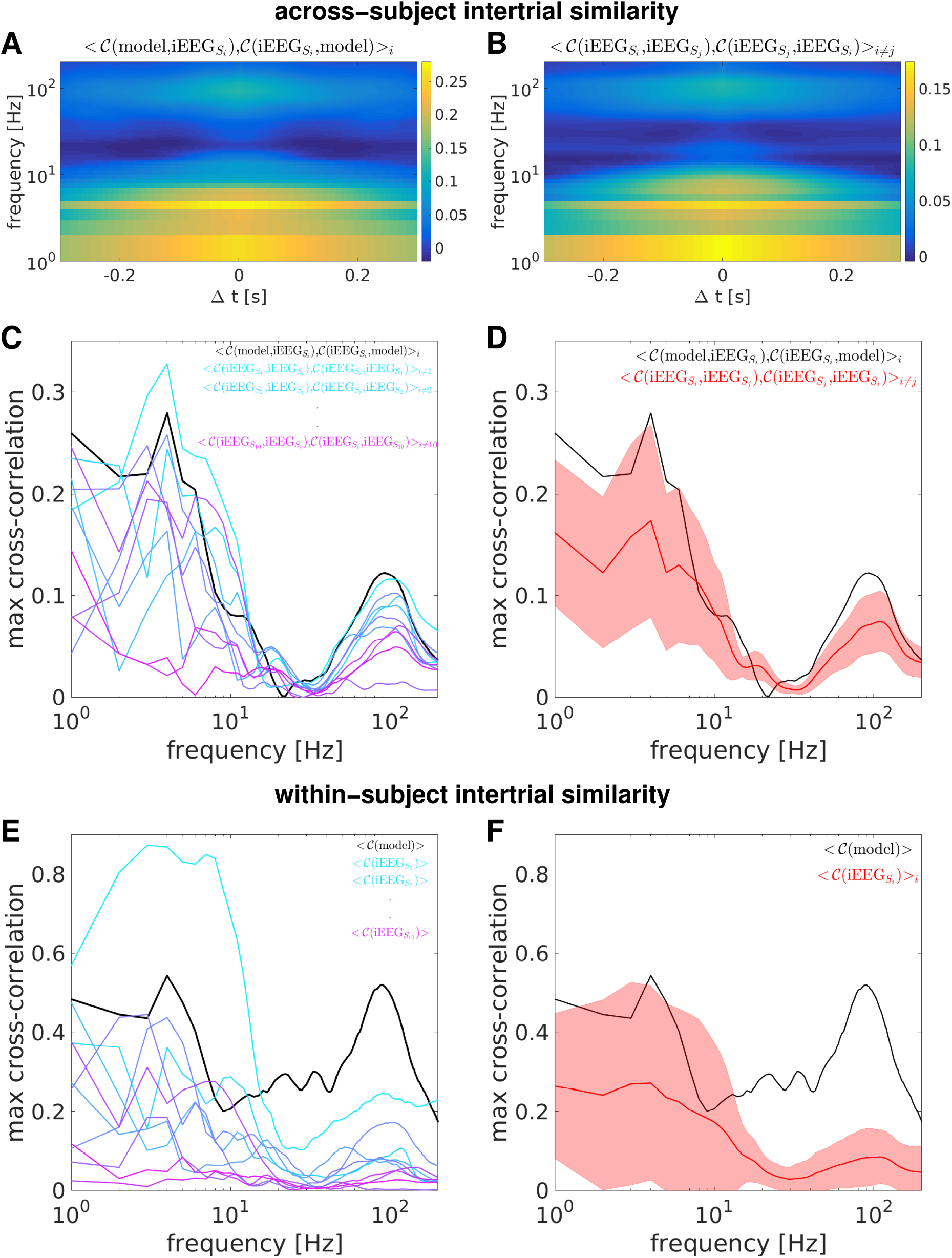
The spectrally-resolved profile of similarity between the model and the iEEG dataset is comparable to inter-subject similarity. A: Average spectrally-resolved cross-correlation between the model and the iEEG dataset. B: Average spectrally-resolved cross-correlation between pairs of iEEG subjects. C: The spectrally-resolved similarity between the model and the iEEG dataset, obtained by taking the maximum cross-correlation for each frequency from the cross-correlation plots shown in A), is shown in black. Corresponding spectrally-resolved similarity curves between each iEEG subject and the remaining iEEG dataset are shown with colored lines. D: As in C), but the spectrally-resolved similarity curves between each iEEG subject and the remaining iEEG dataset have been averaged across subjects (red line). The red shading indicates the SD across subjects. Both the black and the red curves are bimodal, with high similarity in low (delta-theta) and high (gamma - high-gamma) frequencies, and low similarity for intermediate frequencies (alpha-beta). E: as in C), but lines indicate within-subject intertrial similarity for each subject (colored lines) and for the model (black line). F: as in D), for within-subject intertrial similarities.

We then assessed how this similarity relates to the average similarity across iEEG subjects (inter-subject variability) by running the same analysis considering every pair of iEEG subjects (Fig. 3B). These analyses revealed that both the model - iEEG similarity and the iEEG - iEEG similarity profiles exhibit a bimodal shape, with high values at low (delta-theta) and high (gamma - high-gamma) frequencies, and low values for intermediate frequencies (in the alpha-beta range) (Fig. 3C,D).

Finally, we assessed the within-subject intertrial similarity in the same way as described above for the inter-subject similarity, but computing the average cross-correlation between the 10log_10_ power trajectories for each pair of trials 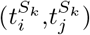, where *i* ≠ *j*, for each subject S_*k*_ as well as for the model (Fig. 3E,F). The spectral profile of within-subject intertrial similarity was qualitatively similar to the spectral profile of inter-subject similarity: it also exhibited the same bimodal shape, albeit similarity values tended to be higher for the within-subject similarity than for the inter-subject similarity, especially in the delta-theta range. The within-subject intertrial similarity in the gamma and high-gamma range was higher in the model than in the iEEG population, suggesting that the model does not accurately reproduce the iEEG trial-by-trial variability in this frequency range.

### Stimulus encoding occurs preferentially at delta-theta and gamma frequencies

To assess whether the speech-induced band-limited power and phase modulations described in the previous section convey information about speech tokens, we performed some multivariate classification analyses on the CWT representation of individual trail voltage trajectories, as described in subsection “Multivariate classification analyses”. In particular, we trained and tested multivariate classifiers using single-trial spectro-temporal representations of iEEG signals corresponding to either baseline activity, or activity corresponding to neural processing of a speech token. We considered all possible within-phrase binary decoding, as well as the binary decoding analyses between corresponding speech tokens in P1 and P2. We further grouped decoding analyses into three categories: detection (speech token vs. baseline), easy discrimination (between different speech tokens, or the same speech token immediately following continuous speech or occurring after a pause), and hard discrimination (between the same speech token occurring at sentence onset in the two phrases).

When considering a broad range of frequency values (between 4 and 200 Hz) simultaneously as multidimensional inputs to the classifiers, every selected channel displayed high and strongly significant decoding accuracies for most binary classifications, which in some case attained perfect discriminability (Fig. S5). Conversely, decoding accuracy in the “control” decoding analysis discriminating baseline activity in the two phrases was low and never significant, apart from one channel where decoding accuracy for phase was weakly significant (just above the p=0.01 significance threshold, uncorrected, Fig. S5). For many of the selected channels, decoding accuracy was high and strongly significant also for the hard discrimination analysis, albeit generally lower than in detection or easy discrimination analyses.

When comparing decoding accuracies using power and phase values, we observed that phase decoding generally yields higher accuracy for every channel and in most of the decoding analyses considered (Fig. 4A-B, *p*=2.47 − 10^−07^, two-sided Wilcoxon signed-rank test). The few exceptions comprised exclusively detection decoding analyses, that is, differentiations between speech tokens and baseline activity. In fact, phase and power did not yield statistically different decoding accuracies when the test was restricted to detection decoding analyses (*p*=0.35), while phase outperformed power for both easy and hard discrimination decoding analyses (*p*=1.49 − 10^−08^ and *p*=0.002, respectively). The advantage of phase over power increased with discrimination difficulty, with the effect size (Cohen’s *d*) being 0.79 for easy discrimination and 1.21 for hard discrimination decoding. In particular, phase yielded strongly significant (p*<*0.001) decoding results in 9/10 subjects for the hard discrimination, while power did so only in 4/10 subjects. Primacy of phase information over power information, especially for distinguishing between activity patterns corresponding to different sensory stimuli, is in accordance with several prior reports in auditory, visual and tactile modalities (Luo and Poeppel, 2007; Howard and Poeppel, 2010; Ng et al., 2013; Gross et al., 2013; Schyns et al., 2011; Ronconi et al., 2017; Wang et al., 2018; Baumgarten et al., 2015), in motor signals (Hammer et al., 2013), as well as in more complex paradigms involving working memory and decision making (Rizzuto et al., 2003; Lopour et al., 2013). In our dataset and for the binary decoding analyses we considered, the combination of power and phase did not convey additional information beyond the one conveyed by the most informative between power and phase (Fig. 4C).

**Figure 4.**
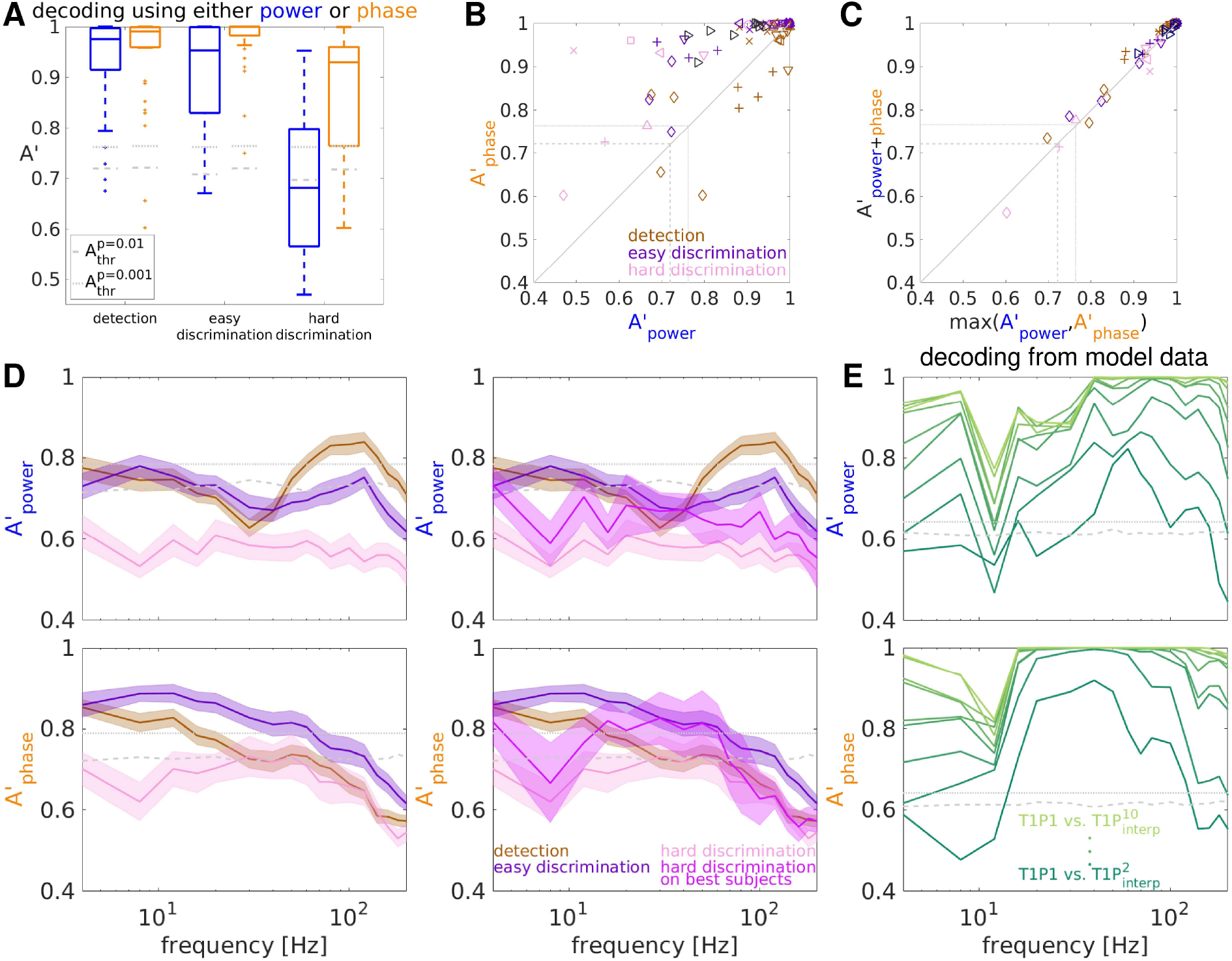
Detection and discrimination of speech tokens from iEEG data. A: Decoding accuracies A’ for three decoding categories using either power (blue bars) or phase (orange bars) values are shown as boxplots: central line indicates the median, box outlines show 25th and 75th percentiles, whiskers extend to the most extreme data points not considered outliers (i.e. up to 1.5 × the interquartile range), and outliers are plotted individually. Decoding accuracies for each decoding analysis are shown for each selected channel separately in Fig. S5. B: Decoding accuracies using phase are plotted against the corresponding decoding accuracies using power for each subject and each classification considered. Colors code for decoding analysis type, as indicated in the legend, while symbol type codes for subject identity as indicated in Fig. S5. Gray dash and dotted lines indicate maximum (across subjects and decoding types) significance thresholds at p=0.01 and 0.001, respectively, obtained by permutation. Apart from some detection decoding analyses, phase is more informative than power. C: Decoding accuracies using power and phase combined are plotted against the corresponding maximum decoding accuracies using either power or phase separately, for each subject and each classification considered. The combination of power and phase does not seem to convey additional information beyond the one conveyed by the most informative variable between power and phase. D: Spectrally-resolved decoding accuracies using either power (top) or phase (bottom) values, for the three decoding types considered. Power discriminability for detection and easy discrimination exhibits a bimodal profile, with high discriminability for low (delta-theta) and high (gamma - high-gamma) frequency values, and low discriminability for intermediate (alpha-beta) frequencies. Right panels also include an additional trace indicating the average hard discrimination decoding accuracy over the four subjects that exhibit significant hard discrimination decoding accuracy across time and frequency at p*<*0.001. Phase discriminability for detection and easy discrimination exhibits a low-pass profile in the group average, but a bimodal profile for hard discrimination is unveiled when averaging only over the four subjects that exhibit highly significant hard discrimination decoding accuracy across time and frequency. Spectrally-resolved decoding accuracies for each decoding analysis are shown for each selected channel separately in Fig. S6. E: Spectrally-resolved speech discrimination decoding in the model. Decoding T1P1 vs. 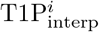 using either patterns of power (top) or phase (bottom) values. Lighter shades of green indicate greater distance from P1. Gray dash and dotted lines indicate significance thresholds at p=0.01 and 0.001, respectively, obtained by permutation.

Next, we proceeded to a spectrally-resolved evaluation of information content by assessing decoding accuracies for a set of frequency values between 4 and 200 Hz. When decoding using power values, the profile of decoding accuracies for detection and easy discrimination parallels our previous results on speech-induced power changes: we observed a timescale separation between high decoding accuracies at low (delta-theta) and high (gamma - high-gamma) frequencies, with low and non-significant decoding accuracies for intermediate (alpha-theta) frequency values (Fig. 4D, top left panel). Conversely, when decoding using phase values, the highest decoding accuracies were observed for low frequency values in the delta, theta and low-alpha range, and decreased for higher frequencies (Fig. 4D, bottom left panel). However, a bimodal profile with enhanced discriminability at low and high frequencies was also observed in some individual subject (Fig. S6). Interestingly, such a bimodal profile was also observed in the hard discrimination decoding when considering only the four subjects that exhibited significant hard discrimination decoding accuracy across time and frequency at p*<*0.001. This suggests that, while a broad frequency range might be informative enough for easy discriminations, hard discriminations might unveil the bimodal spectral profile of information content also for discriminations based on phase values.

Variability across trials in our iEEG data is expected to arise from multiple sources: stochastic opening and closing of ionic channels, stochastic synaptic vesicle release, as well as fluctuations in global physiological state which could be related to variations in attention, arousal, mind wandering, and other factors. Our rate model, albeit endowed with a noise term (which represents mostly microscopic stochasticity in the local network), is not expected to fully capture the extent and complexity of the inter-trial variability in the iEEG dataset. In fact, the rate model yields perfect decoding accuracy in all the detection and discrimination decoding analyses presented above.

However, when the model is tested with a challenging discrimination analysis (i.e., discriminating T1P1 vs. 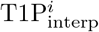, where the latter indicates a stimulus which has been obtained by interpolating between P1 and P2, see Methods for details), it also exhibits a bimodal profile of decoding accuracy as a function of frequency (Fig. 4E). Hence, this profile can be explained as a consequence of the synaptic interactions between excitatory and inhibitory neural populations at theta and gamma timescales, as captured in our model.

The analyses presented so far do not inform on whether the information conveyed by any frequency is specific to that frequency, or whether it generalizes to other frequencies. We estimated the degree of specificity vs. generalization in the read-out of the information conveyed by patterns of either power or phase values using a decoding approach where training and test patterns correspond to different frequency values. When testing speech detection, i.e. distinguishing between neural activity during speech vs. baseline, the information conveyed in patterns of power values largely generalizes across low and high frequency values, but not in intermediate frequency values corresponding to the alpha-beta range (Fig. 5A, left panel). This result is consistent with our characterization of spectral power changes in response to speech stimulation (Fig. 1), and suggests that brain activity corresponding to speech can be distinguished from baseline simply on the basis of a power increase in relatively broad spectral ranges corresponding to low (delta-theta) or high (gamma - high-gamma) frequencies.

**Figure 5.**
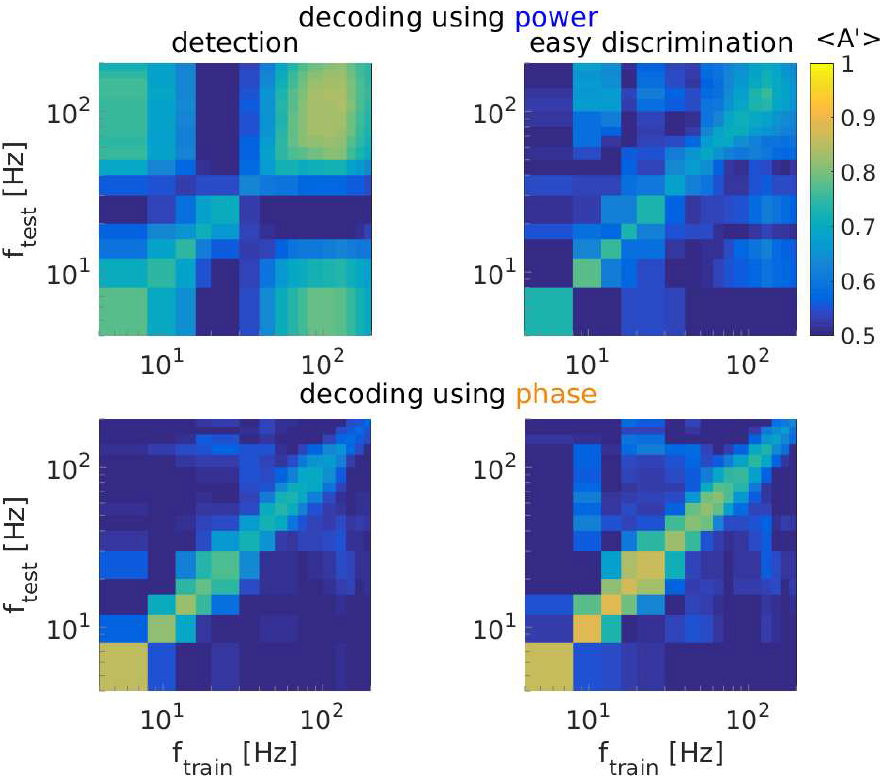
Specificity vs. generalization in the information conveyed by different frequency bands. A: Decoding accuracies A’ averaged across subjects and decoding types when training with patterns of power values extracted from frequency *f*_train_ and testing with patterns of power values extracted from frequency *f*_test_. B: As in A), but considering phase instead of power.

Conversely, the information conveyed by different frequency bands is more specific when considering speech discrimination analyses (distinguishing between neural activity corresponding to different speech segments) (Fig. 5A, right panel), suggesting that the patterns of power values at different frequency bands convey different information about the precise speech segment being heard.

When considering phase decoding, either for speech detection or discrimination, the matrices of decoding accuracies for pairs of (*f*_train_,*f*_test_) only exhibit high values for pairs of frequencies close to the diagonal: decoders tested on frequencies that are sufficiently different from those that are used for training perform poorly, which indicates that speech information is encoded in frequency-specific phase patterns (Fig. 5B).

To summarize, these analyses revealed that patterns of power values convey information for speech detection that largely generalizes across frequencies. Conversely, speech discrimination relies on precise spectro-temporal patterns that are largely frequency-specific. The specificity of information across frequency values is especially conspicuous when considering phase as the relevant information-bearing spectral quantity, which we demonstrated to convey more information than power for speech discrimination (Fig. 4A,B).

### Model-derived relationships between spectral features and stimulus encoding

The relationships between spectral features of neural activity and its stimulus encoding properties is a topic of great interest in sensory neuroscience (e.g., (Belitski et al., 2008; Belitski et al., 2010; Tsuchiya et al., 2008; Schyns et al., 2011; Gross et al., 2013; Ronconi et al., 2017)). The development of a neuronal network model that exhibits remarkable correspondence to iEEG data in terms of both spectral properties and stimulus encoding enabled us to formulate an approach for relating these two aspects of neural activity, which have hitherto been mostly investigated separately.

In particular, we generated family of models by varying some key parameters from their values in the canonical model shown in previous sections. This enabled us to generate “families” of models that are identical, apart from the specific parameter values being varied. The parameters that have been varied are the bias current injected in either Ge or Te units 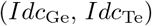), and the strength of recurrent excitatory connections in either the fast or the slow subnetwork (*J*_GeGe_, *J*_TeTe_). A linear system analysis indicates that these parameters affect the network dynamics in different ways: an increase in the bias current tends to linearize the dynamics of the corresponding rate units, while an increase in the recurrent excitatory coupling tends to shift the eigenvalues of the linearized system towards the right, hence increasing the oscillatory character of the subnetwork at its natural frequency. We conducted two sets of simulations: in one set, we varied the bias current and the excitatory self-connection strength in the gamma subnetwork in a two-dimensional interval, while keeping all other parameters at their canonical values (as reported in Table S3); in the other set, we varied the bias current and the excitatory self-connection strength in the theta subnetwork in a two-dimensional interval, while keeping all other parameters at their canonical values. Each individual model has been evaluated in terms of both spectral features (in particular, spectral power in the delta-theta and gamma bands) and stimulus encoding properties (considering the challenging discrimination decoding analysis T1P1 vs. 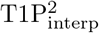. This enabled us to assess the relationships between spectral features and stimulus encoding in a much larger population than our iEEG dataset. In addition, stimulus encoding could be estimated using a higher number of trials than those available in our iEEG datasets (100 per stimulus type as opposed to 30-50 in iEEG subjects), where trial numbers are limited by experimental and clinical constraints, hence yielding more accurate estimates.

These simulations and analyses showed that the information conveyed by patterns of phase values is robustly higher and more stable in the face of parameter variations in comparison to that conveyed by patterns of power values (Fig. S7). They also showed that the strength of recurrent excitatory connections in either the fast or the slow subnetwork (*J*_GeGe_ or *J*_TeTe_) is the parameter that has the largest effect on the decoding performance. Very high values of the recurrent excitatory connection strength impair decoding accuracy, especially when coupled with strong bias currents to the excitatory units. In particular, decoding accuracy using phase is always above 0.95, except in the network with the highest *J*_EE_ and *Idc*_E_ tested, where it decreases to 0.947 (when varying gamma subnetwork parameters) and 0.906 (when varying theta subnetwork parameters). For those values, decoding using power yields an average A’ below 0.79 in both cases. This impairment in coding properties likely derives from an excess of baseline synchronization in the corresponding frequency, and, perhaps more critically, a strong oscillatory responsiveness to inputs impairing selective (hence informative) modulations by the speech input: similar speech segments elicit very similar oscillatory patterns (especially in terms of power, but also in terms of phase), close to saturation. It is worth remarking that this decrease in decoding accuracy is more pronounced when considering patterns of power values, suggesting that, during regimes of strong baseline oscillations and strong oscillatory responsiveness, power modulations induced by temporally patterned inputs are poorly reflected in the network output, whereas the precise phase of neural activity still bears substantial information about the stimulus.

Overall, this analysis shows that the model network encodes speech input with a consistent fidelity across a broad range of parameters, with phase consistently bearing more information than power. Stimulus encoding only deteriorates if the self-excitation strength and the depolarizing excitatory drive in a subnetwork are increased to very high levels, close to dynamic instability, yielding a regime which could be likened to a pathological synchronization, i.e., a seizure-like state (Fig. S7 and S8).

## Discussion

We recorded intracortical responses to pure tones and short sentences from early auditory cortical areas. Based on a combination of anatomical (proximity to Heschls gyri) and functional (maximum ERP amplitude in response to pure tones) criteria, we selected electrode contacts capturing the neural activity of the primary or secondary auditory cortex and characterized their activity in terms of spectral power and phase-locking during the pre-stimulus baseline period and during speech perception. Pre-stimulus activity was characterized by a prominent power peak in the beta range. Speech caused the disappearance of the beta peak and an increase in power in two separate frequency bands, one located in the low frequency range, around the classically defined delta and theta ranges (~ 1-10 Hz), and another one in a high frequency range, corresponding to gamma and high-gamma activity (50-150 Hz). Conversely, phase-locking exhibited a low-pass profile at the group level, even if a bimodal profile could be detected in some subjects (Fig. 1).

We assessed the information encoding properties of iEEG activity by performing decoding analyses probing two aspects of speech perception, detection (distinguishing between neural activity corresponding to a speech segment versus pre-stimulus baseline) and discrimination (distinguishing between neural activity corresponding to different speech tokens). We considered either patterns of power values or patterns of phase values, thereby evaluating the encoding properties of these two distinct aspects of rhythmic neural activity, as well as patterns obtained by concatenating power and phase values. In accordance with previous studies, we demonstrated that phase information generally outperforms power information, with the few exceptions belonging exclusively to detection decoding analyses. The advantage of phase over power increased with discrimination difficulty, with only phase yielding strongly significant decoding results in the group average for the hard discrimination. Combining power and phase information did not improve decoding accuracy beyond the level obtained by the most informative of the two variables.

Consistent with power spectral change analyses, decoding analyses showed that information about the presence of speech (detection) conveyed by patterns of power values exhibited a bimodal profile, with a peak in the low (delta-theta) frequency range and another peak in the high (gamma - high-gamma) frequency range. In addition, the information for speech discrimination conveyed by patterns of power values also exhibited such a bimodal profile (Fig. 4D), with the exception of the hard discrimination decoding in some subjects, which did not exceed the significance threshold at any frequency (Fig. S6). Conversely, the information conveyed by phase patterns exhibited a low-pass profile at the group level, even if a bimodal profile with enhanced discriminability at low and high frequencies was also observed in some individual subject (Fig. S6).

We then addressed whether the neural dynamics reflected in iEEG data could be reproduced by a simple model that assumes that theta and gamma-scale neural activity is underpinned by two interconnected subnetworks, each producing pseudo-rhythmic behavior at distinct timescales. The choice of this architecture is motivated by our own data and is consistent with previous experimental work (Lakatos et al., 2005) and theoretical proposals (Giraud and Poeppel, 2012; Hyafil et al., 2015a; Hyafil et al., 2015b) suggesting that such subnetworks could colocalize in superficial layers of the auditory cortex (see also (Traub et al., 1996; White et al., 2000; Gloveli et al., 2005) for related work in the hippocampus). We developed a population rate model comprised of two subnetworks (each composed of a pair of self and mutually connected excitatory and inhibitory units) with different timescales, one exhibiting fast activity in the gamma - high-gamma range (85.9 Hz), and the other slow activity in the delta-theta range (3.8 Hz), with an inter-subnetwork connectivity pattern implementing a negative feedback loop between the fast and the slow subnetwork.

This simple model exhibits a remarkable similarity to our iEEG channel sample, both in terms of spectral properties at baseline and during speech stimulation (Fig. 2), and in terms of spectrally-resolved information about speech discrimination as measured by decoding analyses (Fig. 4E). These features, and in particular the spectral speech discrimination accuracy profile, do not require fine-tuning of the model parameters, with speech discrimination deteriorating only when the self-excitation strength and the depolarizing excitatory drive in a subnetwork are increased to non-physiological levels, leading to a hyper-synchronized, seizure-like, state.

### Dual timescale processing in auditory cortex

Altogether, our findings are consistent with a dual timescale view of sensory processing in auditory cortex (Giraud and Poeppel, 2012; Teng et al., 2016; Teng et al., 2017), where stimulus encoding is not uniform across frequency bands, nor does it merely reflect the spectral content of the input, but it is instead realized in distinct frequency bands corresponding to low (typically, delta and theta) and high (gamma and high-gamma) frequency bands. Our approach went beyond classical univariate assessment of speech-induced power changes by investigating the information content that each frequency band conveys about the stimulus using spectrally-resolved decoding. We observed that information about the presence of a stimulus (detection decoding), as well as information about stimulus identity (discrimination decoding), were both predominantly conveyed in low (delta-theta) and high (gamma, high-gamma) frequency bands when we considered power and mostly by low frequencies when phase was taken into account, with high frequencies being also informative in some subjects. Interestingly, such a dual timescale profile of neural coding has also been reported in the visual system (Kayser and König, 2004; Belitski et al., 2008; Belitski et al., 2010; Lewis et al., 2016b), suggesting that it might be a general organizational principle of sensory neural processing.

Intermediate frequency bands (alpha and beta) also showed power responses to stimulation (the classical beta power reduction), but this phenomenon was not accompanied by informational activity that could enable reliable stimulus detection or discrimination at the single-trial level using patterns of power values. In addition, activity in these frequency bands was highly variable across individuals, suggesting that activity in this frequency bands is not as strongly driven by sensory stimulation as activity in lower (delta-theta) and higher (gamma - high-gamma) frequencies. Hence, our results lend additional support to the hypothesis that neural activity in the alpha and beta bands might predominantly reflect processes that are not directly involved in the bottom-up encoding of sensory inputs. The low-beta frequency range is indeed thought to be involved in several cognitive aspects of top-down control (Fontolan et al., 2014; Bastos et al., 2015; Park et al., 2015). In the context of speech perception, alpha-beta activity might be predominantly involved with attention (Strauß et al., 2014; Wöstmann et al., 2016; Wöstmann et al., 2017; Dimitrijevic et al., 2017), working memory (Obleser et al., 2012; Wilsch et al., 2015; Wilsch and Obleser, 2016), listening effort (Obleser et al., 2012; Wöstmann et al., 2015; Miles et al., 2017), intelligibility (Obleser and Weisz, 2012; Becker et al., 2013; Pefkou et al., 2017), as well as semantic (Wöstmann et al., 2015), syntactic (Bastiaansen et al., 2010; Bastiaansen and Hagoort, 2015; Lewis et al., 2016a) or temporal predictions (Arnal and Giraud, 2012; Arnal et al., 2015; Morillon and Baillet, 2017); these cognitive aspects (experimentally dissociable only to a partial extent) are consistent with a mostly top-down role.

### Circuit motifs

Substantial progress in our mechanistic understanding of neuronal network function, and on the relationship between structure and activity in brain networks, came from the exploration of circuit motifs and of how they can implement basic neural computations (Womelsdorf et al., 2014; Silver, 2010). In this work we focused on a circuit motif at the inter-subnetwork level, namely the negative feedback loop between the fast and the slow subnetworks implemented by the Ge to Te and the Te to Gi connections. We showed how this inter-subnetwork connectivity configuration results in a remarkable correspondence to iEEG data in terms of both spectral properties and stimulus encoding.

Additional circuit motifs are also likely to be realized in the neuronal circuitry of early auditory cortical areas, and might further contribute to shaping the spectral responses to stimuli as well as to their encoding. For example, feed-forward inhibition has been described in ascending projections from the thalamo-recipient layer L4 to L2/3 in the mouse primary auditory cortex (Li et al., 2014). In our simulations, we observed that feed-forward inhibition (implemented by conveying the bottom-up speech envelope input to the Gi unit as well, in addition to the Ge unit) can modulate the relative proportion of high- vs. low-frequency power responses to stimulation, with increased feed-forward inhibition resulting in lower low- vs. high-frequency power change ratio. However, our canonical model does not include feed-forward inhibition for the sake of parsimony, as an exhaustive characterization of the effect of feed-forward inhibition in our model is beyond the scope of this work.

### Spectral biases and scale-free spectral profiles

Our iEEG recordings tended to exhibit a scale-free power spectrum profile across a fairly broad frequency range (spectral power decreased as 1/f^*α*^), as commonly observed in recordings from the brain and other complex systems (e.g., (He et al., 2010)). Scale-free behavior was prevalent during periods without auditory stimulation (i.e. pre-stimulus periods, Fig. 1C,E, cyan traces). However, it is worth noting that approximate scale-freeness in the power spectrum profile can coexist with band-limited power changes, here expressed as speech-induced synchronization in the delta-theta and gamma ranges and desynchronization in the alpha-beta range. Rather than being realized by a “rotation” of the power spectrum profile as suggested in (Podvalny et al., 2015), our data indicate that opposing changes in average power across neighboring frequency bands (in particular, between speech-induced beta desynchronization and speech-induced gamma synchronization) result from a speech-induced modification of the power spectrum profile towards an inverted double “S” shape in a broader frequency range, corresponding to delta-theta and gamma activation, and alpha-beta deactivation (Fig. 1C,E, red traces). Our decoding results further confirm the presence of a timescale separation in our dataset, with low (delta-theta) and high (gamma - high-gamma) frequencies yielding high and strongly significant decoding accuracies, while intermediate (alpha-beta) frequencies yielded low, often non-significant, decoding performance.

Interestingly, the frequency ranges that exhibited speech-induced activation and speech-induced deactivation varied across subjects, with the transition between the low-frequency activation and the mid-frequency deactivation varying between 7 and 22 Hz, and the transition between the mid-frequency deactivation and the high-frequency activation varying between 20 and 40 Hz (Fig. 2F and S3). Band-limited processes that occur with a consistent structure in all subjects, but at different frequency values, are diluted when data is pooled across subjects, resulting in spectral profiles that are closer to scale-free curves (compare the group-averaged power spectra shown in Fig. 2C, black and gray lines, with the single-subject power spectra shown in Fig. 1C and S2). Hence, we expect band-limited processes to be more readily detected with detailed analyses that independently consider individual channels and behavioral conditions, as enabled by high signal-to-noise ratio recording modalities such as intracranial electrophysiology. Conversely, scale-free behavior is more likely to emerge when data is pooled across channels or behavioral conditions, or when a narrower frequency range is considered, e.g. as in (Miller et al., 2009; Podvalny et al., 2015).

### Potential pre-cortical contributions to cortical oscillations

Oscillatory signatures recorded in the cortex partly originates in the thalamus or in thalamocortical interactions (Steriade et al., 1993; Hughes et al., 2004; Crunelli and Hughes, 2010; Saleem et al., 2017; Li et al., 2017). However, cortical microcircuitry also has the potential to general rhythmic oscillations in multiple frequency bands and might do so, at least in some contextual and neuromodulatory conditions (Buzsáki and Chrobak, 1995; Buzsáki and Draguhn, 2004; Buzsaki, 2006; Wang, 2010). In particular, prominent delta, theta and gamma activity has been localized in superficial layers of the auditory cortex on the basis of current-source density analyses of multi-layered recordings in monkeys (Lakatos et al., 2005) and humans (Halgren et al., 2018).

While the current model implements cortical oscillations as locally generated and assumes a simple, non-oscillatory scheme for pre-cortical stages of auditory processing, the physiological interpretation of each of the model components could be modified to accommodate a role of pre-cortical structures in the generation of rhythmic activity as detected by intracranial EEG in auditory cortex, for example by assuming that theta units could reflect not only local cortical activity but rather the composite activity of neural populations distributed across the cortex and key subcortical structures of the auditory pathway such as the inferior colliculus and the auditory thalamus, and maybe the hippocampus, the cerebellum and the basal ganglia.

### Modeling considerations

Population rate models based on the Wilson-Cowan formalism have been successfully applied to characterize ECoG recordings. In particular, they constitute a simple and suitable description for the 1/f^*α*^ spectral profile of human ECoG activity and its modulation by task engagement (Chaudhuri et al., 2017). Such models have also been successfully applied to the description of feedforward and feedback processing, and their spectral fingerprints, in a large-scale model of the primate visual system (Mejias et al., 2016).

In contrast to previous proposals (e.g., (Loebel et al., 2007; Hyafil et al., 2015a; Stringer et al., 2016; Yarden and Nelken, 2017; Harper et al., 2016; Rahman et al., 2019; Chambers et al., 2019)), our network model does not incorporate a tonotopic structure. This approach is motivated by the psychophysical observation that speech comprehension can be accomplished based mostly on temporal cues (Shannon et al., 1995). Our goal was not to implement a detailed model that incorporated all the known features of the auditory system, but rather to build a minimal model that could reproduce the key features observed in our iEEG datasets, most prominently the presence of discontinuous timescales that characterizes cortical responses to auditory speech stimuli.

In contrast to most previously proposed model of auditory processing (Viemeister and Wakefield, 1991; Moore, 2003; Teng et al., 2016), ours is instantiated at a biophysical level, and relies on specific synaptic interactions between a fast, gamma-scale subnetwork, and a slow, theta-scale subnetwork. As our understanding of auditory processing evolves from phenomenological, abstract models to biophysically grounded implementations, we can develop theories that can be tested not only at the psychophysical level, but also at the level of neural activity patterns. In addition, biophysical models pave the way towards bridging between the neuronal microscale, the iEEG mesoscale and perceptual levels, and bear the potential to relate microscopic anomalies observed in conditions such as autism or dyslexia to corresponding mesoscale anomalies, and, ultimately, to the perceptual deficits in speech and/or reading comprehension that underlie some of their debilitating consequences (e.g., (Wang and Krystal, 2014)).

Biophysical models encapsulate, in an explicit and directly testable format, a set of hypotheses about the underlying neural mechanisms that are deemed sufficient for the emergence of a set of observed features of interest; here, the spectral power features observed at baseline and in response to speech in early auditory cortex, and their information encoding characteristics. From this perspective, the biological features that are not appropriately reproduced by the model are also of great interest, since they highlight the limitations of the current modeling approach and might indicate promising directions for future developments.

With respect to the spectral power profile at baseline and its modulation by speech stimulation, a key feature that our model does not reproduce is the presence of a power peak in the beta range, which is reduced by speech stimulation. In contrast to rhythmic activity at lower (theta, alpha) or higher (gamma) frequencies (Buzsáki and Chrobak, 1995; Wang, 2010), the biophysical mechanisms underlying beta rhythmicity are not well understood. A recent study based on *in vivo* data from humans, monkeys and mice suggested that beta activity could originate from near-simultaneous volleys of excitatory synaptic drive impinging on proximal and distal dendrites of pyramidal neurons (Sherman et al., 2016), which could conceivably originate from the lemniscal and the nonlemniscal thalamus, respectively. Conversely, previous *in vitro* work proposed a mechanism for the emerge of beta rhythmicity, and in particular its slower variant (with spectral peak ~ 15 Hz), based on the concatenation of gamma and fast beta across superficial and deep cortical layers, respectively (Kramer et al., 2008).

Regardless of the mechanistic details of beta activity generation, prominent activity in this frequency band has been associated to a status-quo regime where changes in sensory/cognitive states are not expected (Engel and Fries, 2010) and local information processing and propagation are mechanistically inhibited (Sherman et al., 2016), maybe in favor of a more global operating mode (Kopell et al., 2000; Kopell et al., 2011). Neural activity in the beta range exhibits a prominent top-down directional bias across the neural hierarchy (Buschman and Miller, 2007; van Kerkoerle et al., 2014; Fontolan et al., 2014; Bastos et al., 2015; Bastos et al., 2015; Michalareas et al., 2016), lending strong support to the hypothesis that beta oscillations as recorded in early auditory cortical areas might be largely inherited from higher cortical areas. From this perspective, it is not surprising that our computational model, designed to capture the essential neuronal microcircuitry of the superficial layers of a single cortical area, does not reproduce the beta power spectrum peak at baseline and its reduction upon speech presentation. Future modeling work will address the interplay of bottom-up sensory inputs and top-down expectations or linguistic knowledge by developing a hierarchical neuronal network with spiking neurons and realistic laminar organization (e.g., (Bastos et al., 2012; Lee et al., 2013; Lee et al., 2015)). Importantly, the development of neuronal network models that incorporate linguistic knowledge could pave the way towards the discernment and a mechanistic understanding of the micro- and meso-scale origins of the macroscopic features in brain oscillation power (Obleser and Weisz, 2012; Becker et al., 2013; Pefkou et al., 2017), coherence with speech (Luo and Poeppel, 2007; Peelle and Davis, 2012; Doelling et al., 2014; Kösem and van Wassenhove, 2017), and scale-free amplitude dynamics (Borges et al., 2018) that selectively accompany speech comprehension.

Another feature of interest that our model only reproduce to a partial extent is inter-trial variability, a feature of neural activity that is directly related to information encoding. Variability across trials in our iEEG data is expected to arise from multiple sources: stochastic opening and closing of ionic channels, stochastic synaptic vesicle release, as well as fluctuations in global physiological state which could be related to variations in attention, arousal, mind wandering, and other factors. Our rate model, albeit endowed with a noise term (which represents mostly microscopic stochasticity in the local network), is not expected to fully capture the extent and complexity of the inter-trial variability in the iEEG dataset. Notably, and in spite of its inability to quantitatively match iEEG inter-trial variability, our model network qualitatively reproduces the bimodal spectral profile of decoding accuracy for speech discrimination observed in our iEEG dataset (Fig. 4E), when discrimination is assessed by considering activity patterns corresponding to highly similar speech inputs. Further modeling work could incorporate additional terms with slowly varying dynamics accounting for specific aspects of brain physiology that are expected to affect neural variability, such as sensory adaptation and shifts in attention and arousal (maybe by applying similar strategies as employed in (Goris et al., 2014) for the characterization of variability in spiking activity), hence potentially elucidating the influence of each of these processes on inter-trial variability.

## Conclusion

We showed that neural activity in early auditory cortex in humans, as directly recorded by intracranial EEG, displays separable timescales at low (delta-theta) and high (gamma - high-gamma) frequencies, where speech stimulation increases spectral power and where reproducibility and information for speech detection and discrimination is higher than in the intermediate alpha - beta range. We further showed that a simple rate model comprised of two excitatory/inhibitory subnetworks with different timescales and interconnected with a negative feedback loop between the fast and the slow subnetwork is sufficient to account for a vast majority of the experimental observations. Interestingly, the model failed to reproduce the baseline beta peak and its speech-induced desynchronization, which is consistent with the recent proposals suggesting that beta activity might have different underpinnings, possibly involving dynamic coordination across the cortical hierarchy.

After acknowledging that the main spectral and informational features of neural activity at baseline and in response to speech are well captured by a simple rate model with four neuronal populations, one might feel inclined to believe that the apparent complexity of brain activity could be reduced to a fairly low-dimensional dynamical system. We would suggest a different interpretation, not incompatible with but complementary to the former: if these simple models can capture broad features of the iEEG, these broad features might only convey a degraded version of the information presented at finer resolutions. While this could be enough for a rough reconstruction of the stimulus, it would not allow for re-creating the complex and highly informative phenomenological experience that speech and music can elicit in healthy, awake humans.

It is therefore of interest to assess the extent to which simple models, which embody well-specified hypotheses about the neurobiological substrates (in particular, the connectivity diagram between excitatory and inhibitory populations and the time constants that govern their interactions), can account for various spectral measures of iEEG activity, advancing from relatively gross features such as ERP amplitude, spectral power and phase-locking characteristics, to progressively finer measures related to variability and information encoding and transformation. Hence, we believe that the interaction of advanced data analysis techniques, addressing both dynamical and informational features of brain activity, and computational modeling at multiple levels of resolution can offer a promising toolkit for improving our understanding of the inner workings of the brain, and in particular for clarifying the spatial resolution that maximizes the causal effectiveness of neural activity (Hoel et al., 2013), arguably a necessary prerequisite for a mechanistic understanding of brain function.

## Acknowledgments

This work was funded by a project grant from the Swiss National Fund to ALG (#320030-163040). The computations were performed at the University of Geneva on the Baobab cluster.

