## Supplementary Information for "Converging intracortical signatures of two separated processing timescales in human early auditory cortex"

### Supplementary Tables

| sub ID | age | sex | educational level | language dominance | MRI | seizure focus |
|---|---|---|---|---|---|---|
| S1-RH | 31 | male | 2 | left | normal | right temporal |
| S2-LH | 30 | female | 1 | left | HS | left temporal |
| S3-RH | 36 | female | 3 | left | normal | bilateral temporal |
| S4-RH | 25 | female | 1 | left | normal | bilateral temporal |
| S5-RH | 62 | female | 2 | atypical | normal | right temporo-mesial |
| S6-RH | 42 | male | 2 | left | normal | right temporal |
| S7-RH | 26 | female | 1 | left | normal | bilateral temporo-basal |
| S8-RH | 22 | male | 2 | left | normal | right post-perisylvian |
| S9-LH | 17 | female | 1 | left | FCD type II | left pre-frontal |
| S10-RH | 18 | male | 2 | left | normal | right temporal |

**Table S1: Demographic and clinical information for each subject.** Educational level: 1, no high school graduate; 2, high school graduate; 3, college undergraduate or higher. HS: hippocampal sclerosis; FCD: focal cortical dysplasia.

| sub ID | pure tone frequencies | # of trials, pure tones | # of trials, phrase |
|---|---|---|---|
| S1-RH | Set 2 | 58/48/59/52/54 | 24/29 |
| S2-LH | Set 1 | 79/63/87/76 | 52/58 |
| S3-RH | Set 2 | 67/58/73/62/69 | 52/58 |
| S4-RH | Set 1 | 112/103/109/109 | 30/35 |
| S5-RH | Set 1 | 55/41/54/57 | 32/35 |
| S6-RH | Set 2 | 44/35/43/40/43 | 35/42 |
| S7-RH | Set 3 | 50/50/50/50/50; 50/50/50/50/50 | 51/55 |
| S8-RH | Set 2 | 66/56/73/57/65 | 52/58 |
| S9-LH | Set 2 | 70/62/79/66/72 | 52/58 |
| S10-RH | Set 2 | 57/42/55/59 | 42/48 |

**Table S2: Stimuli and number of trials for each subject.** Set 1: 250 Hz, 750 Hz, 2 KHz, 4 KHz (binaurally presented). Set 2: 250 Hz, 750 Hz, 2 kHz, 3 kHz, 4 kHz (binaurally presented). Set 3: 500Hz, 1 kHz, 2 kHz, 3 kHz, 4 kHz to the right ear; and corresponding stimuli to the left ear.

| Description | Parameter symbol and value |
| --- | --- |
| **Intrinsic parameters** | |
| Population time constant, Ge | $\tau_{\mathrm{Ge}}$=3.5398 ms |
| Population time constant, Gi | $\tau_{\mathrm{Gi}}$=8.8496 ms |
| Population time constant, Te | $\tau_{\mathrm{Te}}$=50 ms |
| Population time constant, Gi | $\tau_{\mathrm{Ti}}$=125 ms |
| **Background input parameters** | |
| DC bias current, Ge | $Idc_{\mathrm{Ge}}$=-2 |
| DC bias current, Gi | $Idc_{\mathrm{Gi}}$=0 |
| DC bias current, Te | $Idc_{\mathrm{Te}}$=4 |
| DC bias current, Gi | $Idc_{\mathrm{Ti}}$=0 |
| Background noise SD, Ge | $\sigma_{\mathrm{Ge}}$=0.1 |
| Background noise SD, Gi | $\sigma_{\mathrm{Gi}}$=0.1 |
| Background noise SD, Te | $\sigma_{\mathrm{Te}}$=0.15 |
| Background noise SD, Gi | $\sigma_{\mathrm{Ti}}$=0.15 |
| **Inter-subnetwork coupling parameters**[1] | |
| Self-coupling, E units | $J_{\mathrm{EE}}$=1.5 |
| Self-coupling, I units | $J_{\mathrm{II}}$=-2.5 |
| Coupling from I to E units | $J_{\mathrm{EI}}$=-3.25 |
| Coupling from E to I units | $J_{\mathrm{IE}}$=3.5 |
| **Intra-subnetwork coupling parameters**[2] | |
| Coupling from Ge to Te | $J_{\mathrm{TeGe}}$=1.5 |
| Coupling from Te to Gi | $J_{\mathrm{GiTe}}$=1.125 |
| **Bottom-up input and synaptic depression parameters** | |
| Bottom-up input strength | $\mathcal{A}$=50 |
| Rate of available neurotransmitter decay | $U$=0.005 ms$^{-1}$ |
| Time constant of available neurotransmitter recovery | $\tau_{\mathrm{D}}$=1000 ms |
| Sigmoid function steepness | $k_{\mathrm{D}}$=1.8 |
| Sigmoid function midpoint | $x_0$=2.2 |

**Table S3: Model parameters.** Parameter descriptions and canonical values used throughout this study, unless otherwise stated. Units are arbitrary, unless stated otherwise. [1] These parameters are the same for both the fast and the slow subnetwork. [2] Only parameter values that differ from zero are listed.

### Supplementary Figures

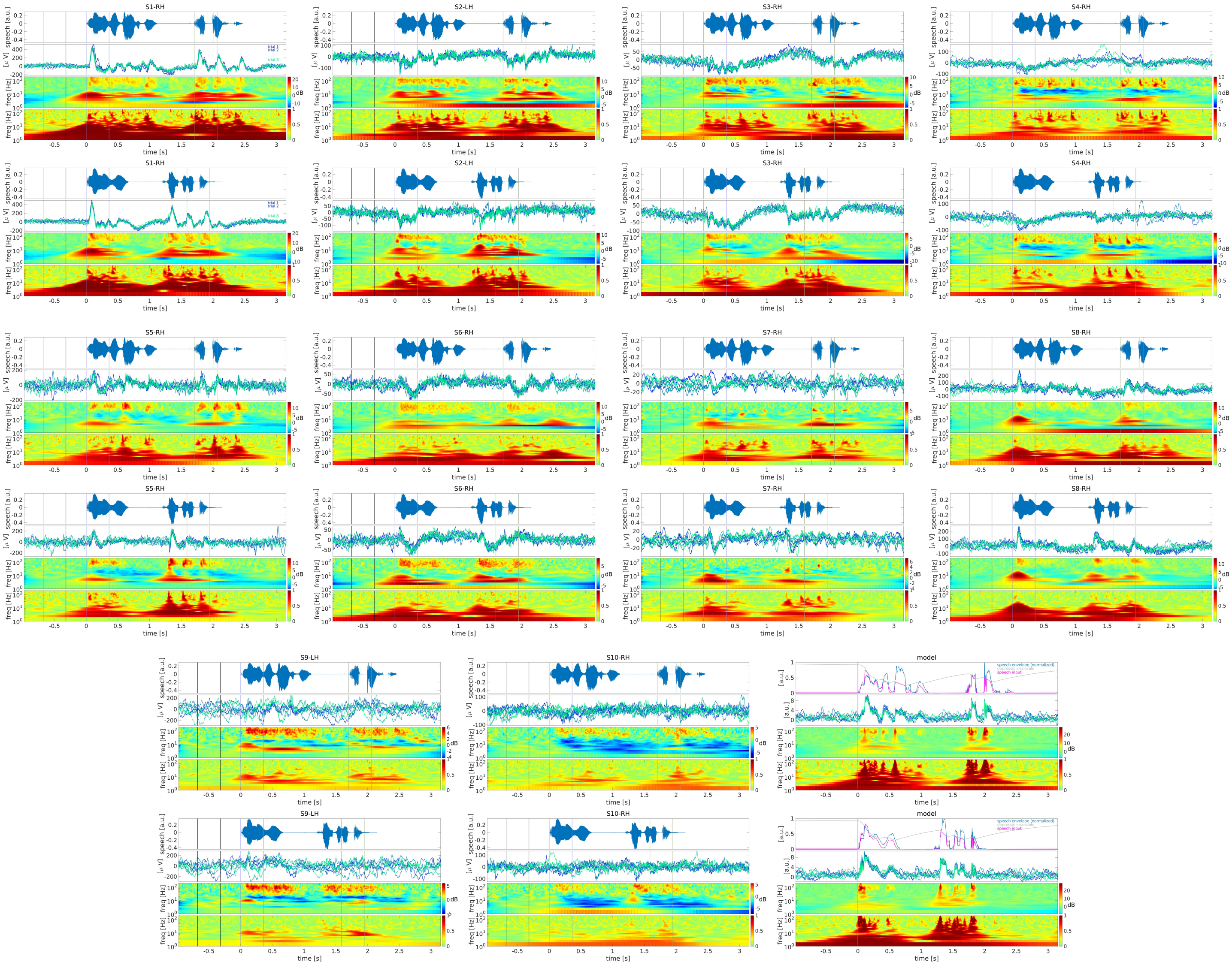

**Figure S1: Voltage trajectories, power and PLV spectrograms.** Same format as in Fig. 1B, for all subjects and for both phrases, as well as for the model.

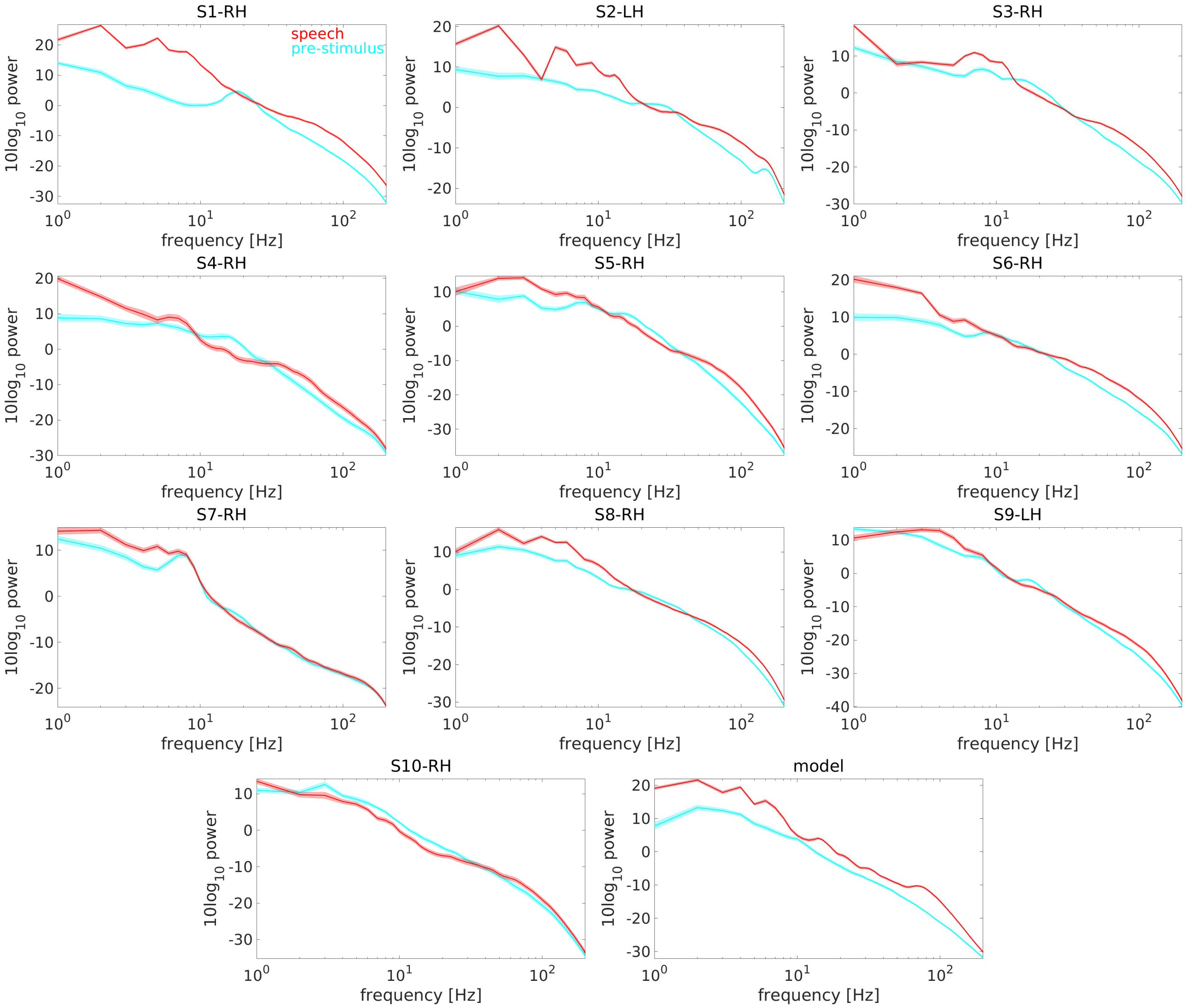


**Figure S2: Power spectra during the pre-stimulus baseline and during speech presentation.** Same format as in Fig. 1C, for all subjects and for the model.

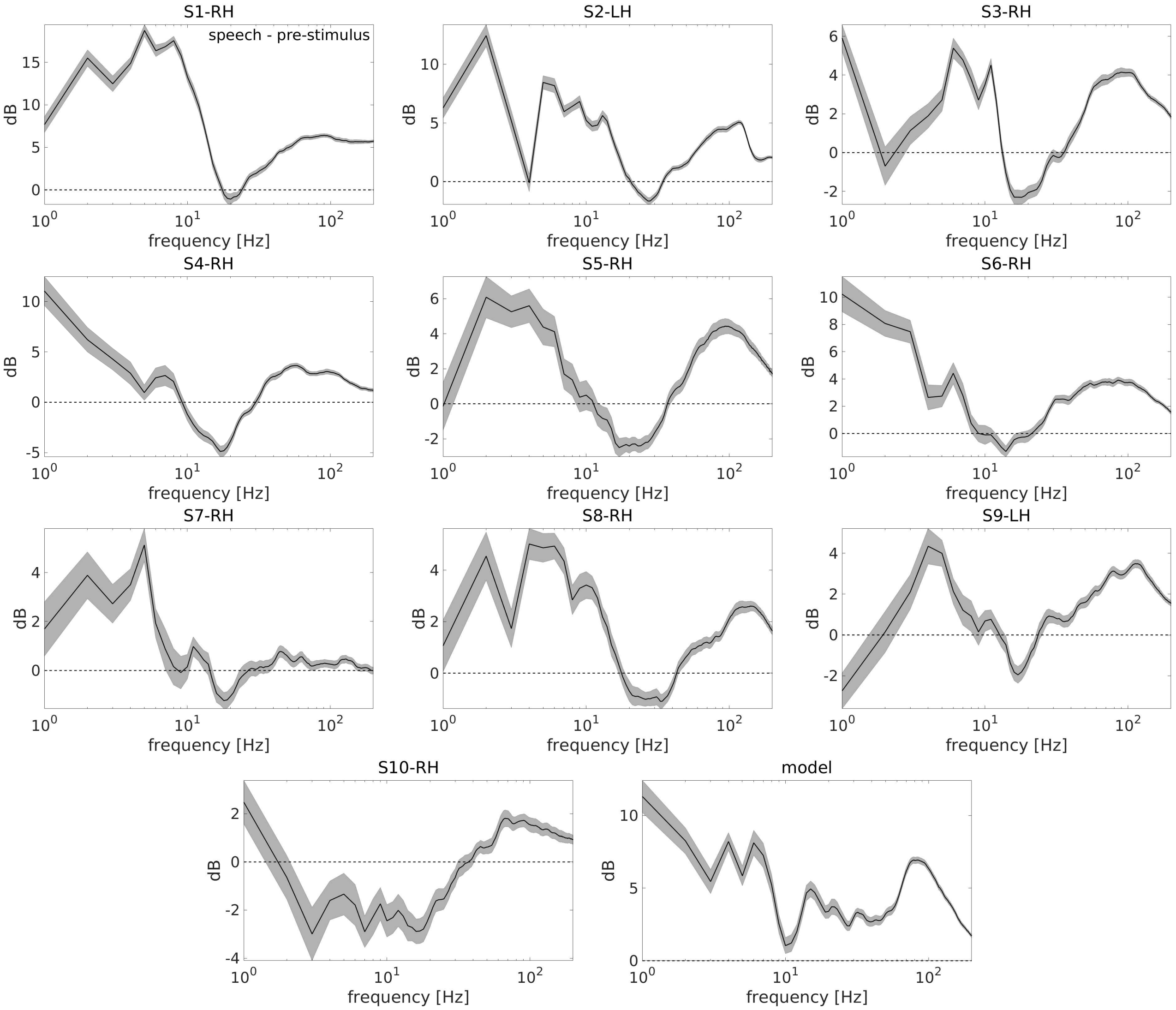


**Figure S3: Power spectrum changes induced by speech presentation.** Same format as in Fig. 1D, for all subjects and for the model.

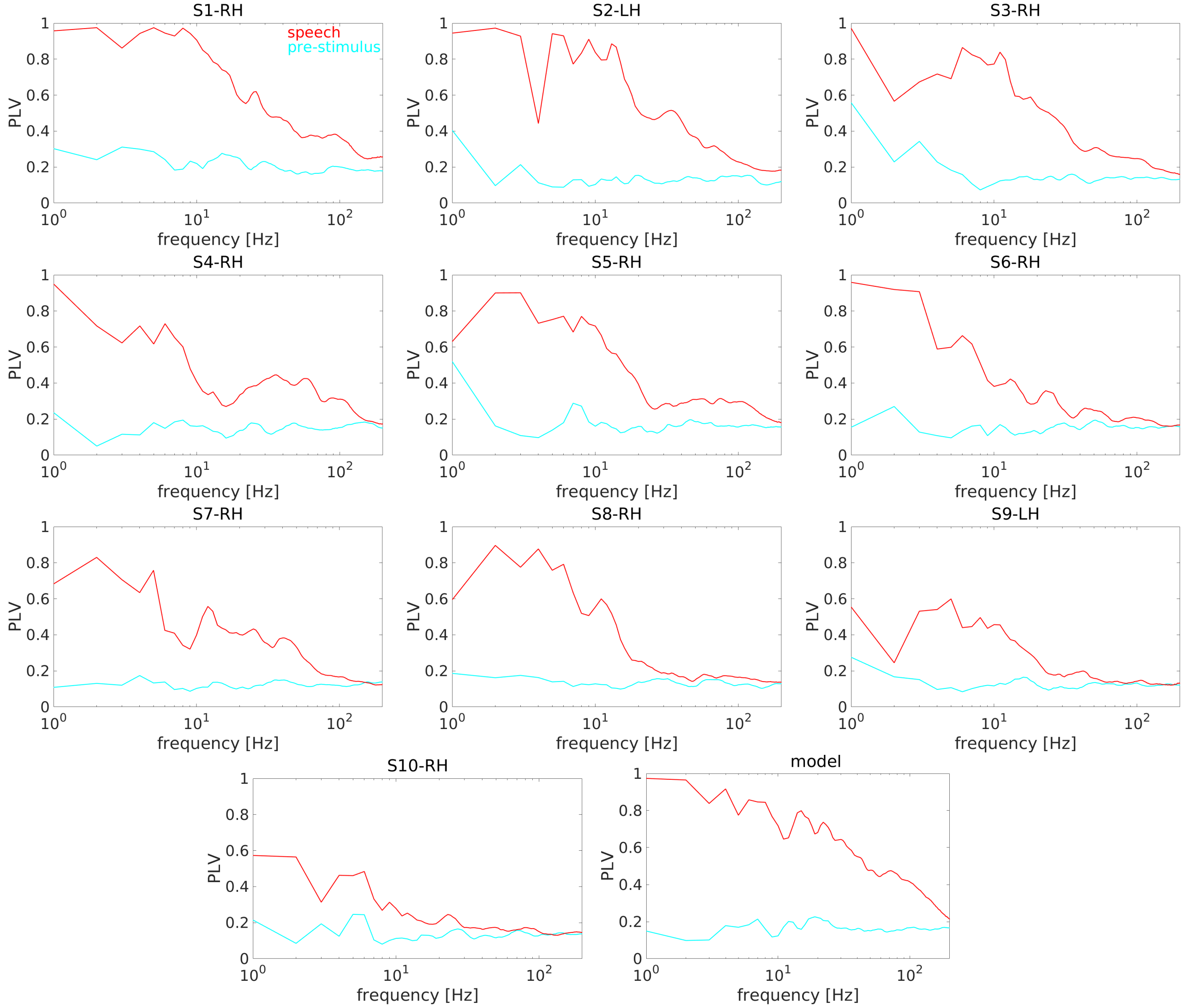


**Figure S4: Phase-locking values during the pre-stimulus baseline and during speech presentation.** Same format as in Fig. 1E, right panel, but each individual channel and the model are shown in separate panels.

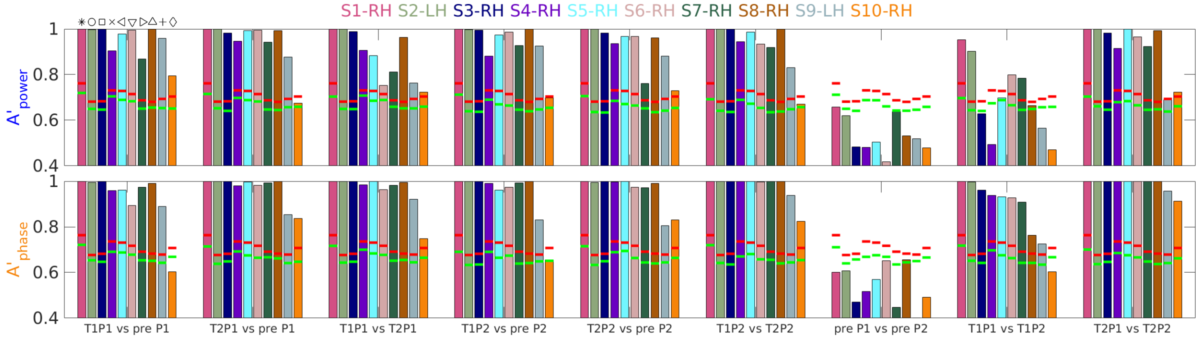


**Figure S5: Decoding accuracy for speech detection and discrimination.** Decoding accuracies A' for each subject and each classification considered. Horizontal lines indicate the p=0.01 and 0.001 significance thresholds, obtained via permutation. Subjects are ordered according to decreasing decoding accuracy in the hard discrimination analysis (T1P1 vs. T1P2) using phase values. Symbols on top of the first group of bars indicate the subject identity code used in Fig. 4B and C.

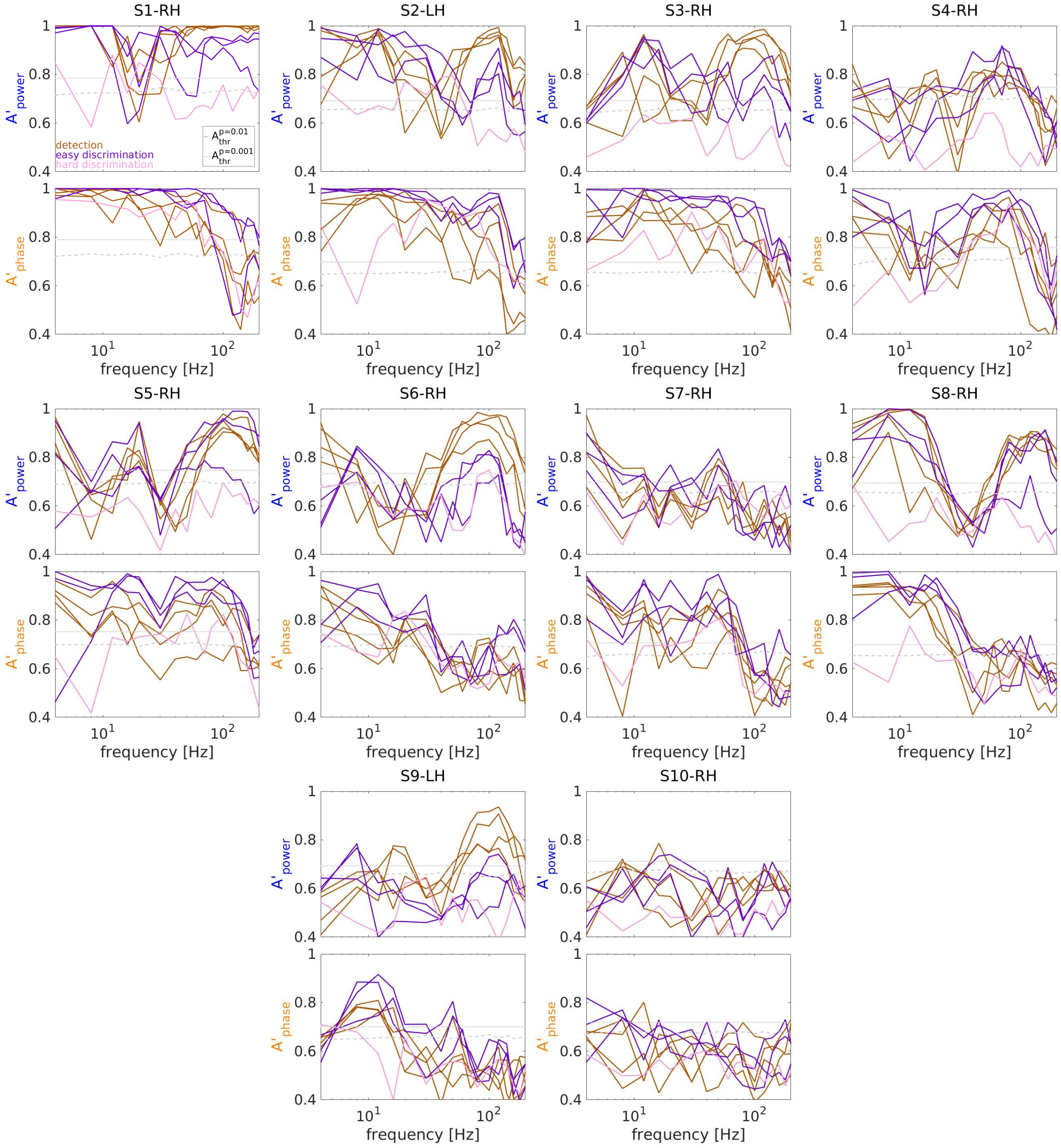


**Figure S6: Spectrally-resolved decoding accuracy for speech detection and discrimination.** Same format as in Fig. 4D, but decoding accuracy for each decoding analysis is shown. Results from each selected channels are shown in separate panels.

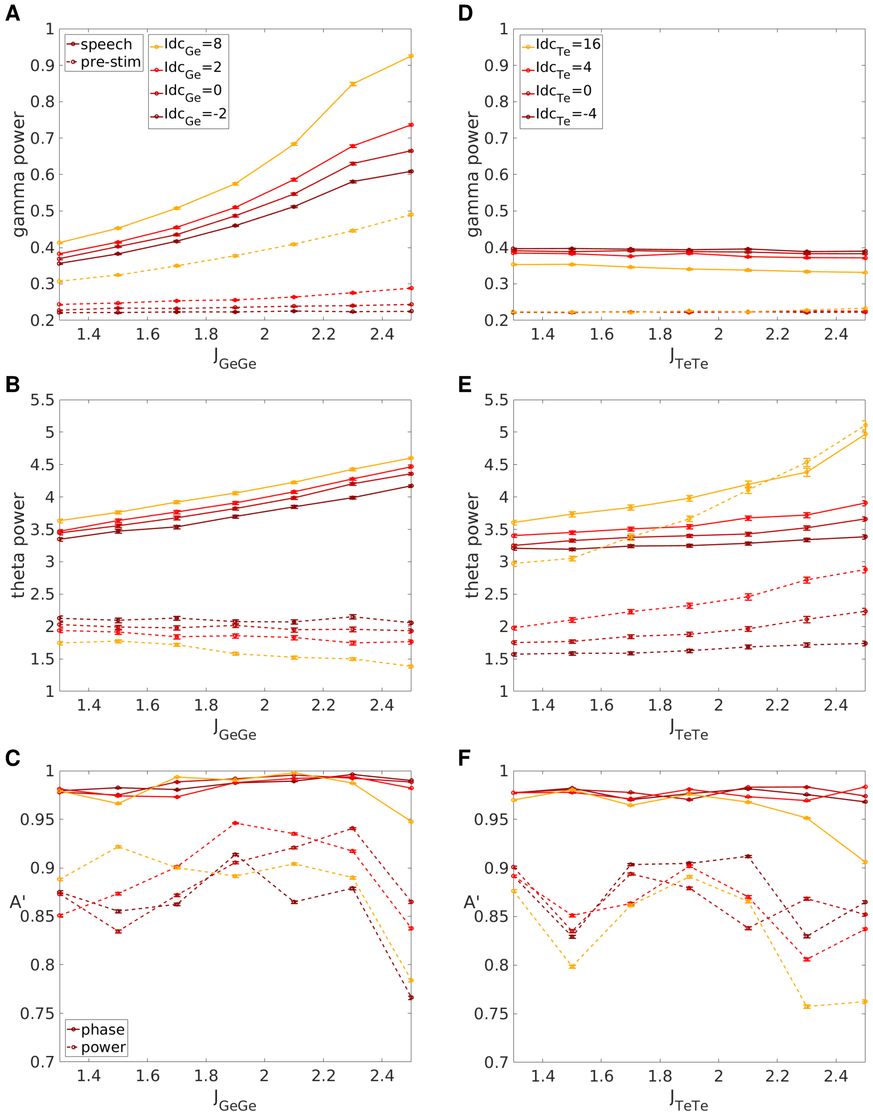


**Figure S7: Dependence of spectral features and stimulus encoding on model parameters.** Effects of varying either gamma subnetwork parameters (left column) or theta subnetwork parameters (right column) on gamma power (top), theta power (middle), and decoding accuracy in the T1P1 vs. T1P$^2_{\text{interp}}$ discrimination analysis (bottom). Power estimates were obtained by computing the continuous wavelet transform of the de-meaned simulated LFP for the baseline ([-1000, 0] ms) and the speech period ([0, 1000] ms) for each trial in response to the P1 stimulus, taking the cubic root of spectral power for each time and frequency point, and averaging over time and corresponding frequency intervals (76-120 Hz for gamma, 3-8 Hz for theta). Error bars indicate s.e.m. across trials (for power estimates) or across cross-validation folds (for decoding accuracy).

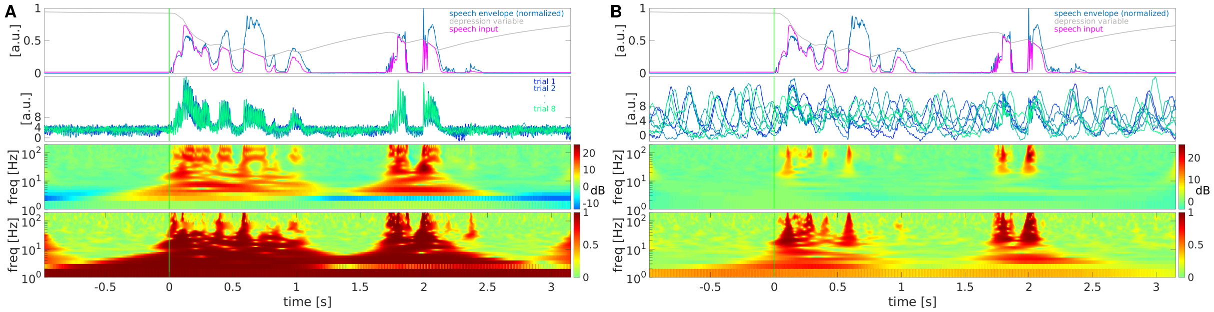


**Figure S8: Strong recurrent excitatory connection and depolarizing excitatory drive saturate oscillatory activity and impair accurate stimulus encoding.** A: Speech input (top), single-trail trajectories (middle-top), power (middle-bottom) and PLV (bottom) spectrograms are shown with the same format as in Fig. 2B, for the network model with the highest values of the recurrent excitatory connection strength $J_{\mathrm{GeGe}}$ and the depolarizing excitatory drive $Idc_{\mathrm{Ge}}$ tested. B: As in A), for the network model with the highest values of the recurrent excitatory connection strength $J_{\mathrm{TeTe}}$ and the depolarizing excitatory drive $Idc_{\mathrm{Te}}$ tested.
